# Differential effect of supercoiling on bacterial transcription in topological domains

**DOI:** 10.1101/2025.01.02.631060

**Authors:** Boaz Goldberg, Nicolás Yehya, Jie Xiao, Sam Meyer

**Affiliations:** Department of Biophysics and Biophysical Chemistry, Johns Hopkins School of Medicine, Baltimore, MD, USA; Université de Lyon, INSA Lyon, Université Claude Bernard Lyon 1, CNRS UMR5240, Laboratoire de Microbiologie, Adaptation et Pathogénie, 69621 Villeurbanne, France

## Abstract

DNA supercoiling (SC), the over- and under-winding of DNA, is generated by transcription as described in the twin-domain model. Conversely, SC also impacts transcription through torsional stress. SC therefore regulates transcription independent of transcription factor binding, the classic protein-based transcription regulatory mechanism, particularly in the context of chromosomal topological domains and the actions of topoisomerases. SC-coupled transcription has inspired several computational models, but a systematic and quantitative assessment of the parameters controlling this complex interplay is missing. There is also a lack of comparison to various experimental results regarding the effects of topological variables such as topoisomerase binding rates and domain size on transcription strength and noise. In this work we developed a quantitative model to describe SC-coupled transcription. We analyzed the effects of topoisomerase activities on the transcription of a single isolated gene with varying promoter strengths and in a topological domain of varying sizes. These simulations revealed qualitatively different roles of the two topoisomerases. Topoisomerase I is specifically required for strongly expressed genes that may be hindered by stalled RNAP, whereas gyrase activity favors the expression of all genes by enhancing transcription initiation and modulating the burstiness of transcription. A new analysis of transcriptomics data in several bacterial species showed that these simulations replicate a global relationship between promoters’ strength and response to variations in topoisomerase activities. Our work demonstrates how SC contributes to differential gene regulation and transcriptional bursting in mechanistic details and unites computational and experimental work.

**Author Summary:** We are interested in understanding how bacteria regulate the expression of their genes. One mechanism by which bacteria can do so is through the over- and under-winding of their DNA, termed DNA supercoiling. Because bacteria can use enzymes called topoisomerases to regulate the supercoiling level of their DNA, these enzymes can serve as gene regulators. It is difficult, however, to understand how topoisomerases act as gene regulators through experiments alone. Currently, there is no method of measuring the supercoiling level along a stretch of DNA over an extended period of time in living cells. We therefore developed a computational model of gene transcription coupled to supercoiling dynamics. Unlike some previously existing models, we consider continuous response curves of topoisomerases’ activities in response to the local supercoiling level and the ability of gyrase to perform several catalytic cycles per binding event. We also perform extensive comparisons between our model and existing gene expression and transcriptomics data, which is essential to ensuring the biological relevance of computational modeling efforts. We replicate and provide detailed explanations for several experimental observations, including the connection between supercoiling and transcriptional bursts and the selective gene expression modulation by topoisomerase inhibition that is based on the promoter strength.

## Introduction

DNA supercoiling (SC), the over- (positive) and under-winding (negative) of double-stranded DNA (dsDNA) relative to its relaxed B-form state, has been proposed as a new means to regulate transcription independent of regulatory protein binding [1]. Negative SC facilitates transcription initiation by driving promoter melting during the formation of the open complex consisting of DNA and RNA polymerase (RNAP) [2]. During transcription elongation, SC-induced torque can stall RNAP [3]. Conversely, transcription also influences SC: an elongating RNAP molecule introduces negative SC upstream and positive SC downstream per the twin-domain model due to the compensatory DNA rotation caused by the rotational drag of the transcription complex [4, 5]. Moreover, SC can diffuse over distances of several kilobases, allowing for cooperative and antagonistic interaction of multiple RNAP molecules within and between genes [6]. Diffusion of SC can be blocked by topological barriers formed by protein binding and/or protein-induced DNA loops, which prevent dsDNA rotation [7] and topologically isolate genes from those on the other side the barriers. Given the coupled interactions between SC and transcription, SC in topological domains may serve as a potent and complex transcriptional regulator independent of protein transcription factors, leading to a DNA-based transcription regulatory network to act upon adjacent genes in long distances.

Cells regulate the average SC level of their chromosomes using topoisomerases. In bacteria, topoisomerase I (TopoI) and gyrase [3] play important roles in maintaining SC homeostasis. TopoI is a type I topoisomerase which relaxes negative SC by introducing transient single-stranded breaks in dsDNA in an ATP-independent manner; gyrase is a type II topoisomerase which hydrolyzes ATP to relax positive SC and introduce negative SC by passing one dsDNA segment through a transient double-stranded break of another [8]. Given the interactions of SC and transcription, it is reasonable to expect that variations in topoisomerase activities induce changes in transcription activities; indeed, prior work on the transcriptomes of several bacterial species has confirmed that inhibition of topoisomerases with antibiotics alters transcription, and that this alteration varies between genes in a manner dependent on growth phase, expression strength, and genomic context [8–18].

While SC may serve as a new, important transcriptional regulator, our understanding is limited by experimental difficulties in measuring the spatiotemporal dynamics of SC along the chromosome and its impact on transcription kinetics. Computational modeling can explore various aspects of SC and its coupling with transcription, help identifying possible underlying mechanisms, and providing testable hypotheses for further experimental development [19–21]. In the past years, we and others have developed multiple models of coupled dynamics between bacterial transcription and SC, either with a mathematical framework [19, 20, 22] or with a computational biophysics focus [12, 17, 21, 23–25]. Some models considered the regulatory effect of SC on transcription initiation [12, 26], elongation [25, 27], or both [17, 21, 23–25]. Some explicitly described the activities of both gyrase and TopoI [12, 17, 23], or the bursting properties of transcription [28]. Additional models relevant to eukaryotic transcription also exist [21, 29].

While these past studies greatly improved our understanding of the relationship between SC and transcription, only a few of them [12, 17, 23, 24, 28] quantitatively compared modeling results with experimental data, mostly focusing on that of Kim *et al*. [6]. Nevertheless, numerous other transcription datasets have been collected since the 1980s, providing valuable, diverse, and complementary quantitative information on the role of topoisomerases and topological constraints on gene expression, including in their native context. These data range from *in vitro* transcription studies involving plasmids prepared at different superhelical densities [30–32], to whole-genome transcriptomics data observed under varying superhelical densities induced by drugs or topoisomerase mutations [9–16, 33], as well as studies of regulatory interactions between neighboring genes in various orientations, based on artificial constructions on the chromosome or plasmids [20, 34, 35]. Systematic integration and interpretation of these data with computational modeling is therefore an important task in the field. To our knowledge, only one computational study has incorporated transcriptomics data in its analysis [12], and only for two bacterial species (*Escherichia coli* and *Dickeya dadantii*). In this study, we aim to integrate computational modeling and experimental transcription data through quantitative simulations of transcription of a single gene isolated in a defined topological domain and a systematic analysis of transcriptomes from many species.

This single-gene approach is justified by the complexity of transcription-SC coupling observed in earlier modeling works (and this work). Due to the highly nonlinear and dynamical features of the process, small quantitative variations in the parameters can lead to qualitatively diverging behaviors. While most models have converged to similar qualitative descriptions of the process, several elements cannot be quantitatively parameterized and still rely on partially arbitrary choices (e.g., the effect of SC on transcription initiation, topoisomerase binding rates, and the size of a topological domain). Another limitation of existing works is the absence of a systematic assessment of how these parameters affect the regulatory behavior of the system, making it unclear if the diversity of reported behaviors results from differences in modeling choices or values of one or several of these factors. In this work, we analyze the effect of three physiologically relevant quantitative parameters, topoisomerase activities, topological domain size, and promoter strength, on the transcription of a single gene and compare it to previously published transcriptomic data of direct relevance (*in vitro* and *in vivo*). As we describe below, this modeling approach produced a remarkably rich and non-intuitive landscape of regulatory behaviors, in particular very different effects for topoisomerase I and gyrase, and a qualitatively different response of weak and strong promoters to topological constraints. Our simulations provide a mechanistic basis for quantitative experimental observations: (1) the characteristic response curve of many promoters to varying SC *in vitro*, with optimal expression observed around physiological negative levels [30–32]; (2) the complex, promoter-dependent regulatory effect of varying topoisomerase activity *in vivo*, as well as topological domain sizes [9–17, 33]; and (3) the emergence of transcriptional bursting in the absence of transcription factor binding [28].

## Methods

### Modeling method overview

Our model broadly follows an approach used in several previous works including ours [12, 17, 23–25] and consists of a stochastic simulation of transcription and one-dimensional (1D) super-helical distribution over time. Briefly, we use a one-step, SC-dependent description of transcription initiation, a stalling threshold on upstream and downstream SC leading to RNAP stalling during elongation, and instantaneous diffusion of supercoils (at the timescale of transcription and over kb-scale distances). The main improvements over existing works are (1) the description of (nonspecific) topoisomerase activity, for which we employ a gradual, realistic response function to local SC, and (2) the introduction of gyrase processivity, based on experimental evidence [36, 37]. While several of these modeling ingredients remain subject to improvements, we verified that the main conclusions of this study are robust to minor changes in choices of functions (e.g., we tested step and logistic functions for topoisomerase binding) and parameters regarding initiation and topoisomerase activities.

We simulate a topological domain containing a single gene of length *L*, with two flanking regions of distance *D* to the domain barriers, giving an overall domain size of 2*D* + L (Fig. 1). We assume that SC diffuses instantaneously on a timescale much shorter than that is relevant to transcription [38], and that both bound RNAP molecules and the ends of the domain serve as topological barriers. These assumptions are in alignment with most previous models [12, 17, 21, 24] and are supported by physical models in the case of twist, which diffuses over domains on the order of 10 kb within 1 ms. While writhe diffuses in a slower and more complicated manner, involving phenomena such as plectoneme hopping, full consideration of this effect would require a three-dimensional (3D) model that would be computationally intractable on timescales relevant to transcriptional regulation [38]. Previously, we have examined finite twist diffusion in transcription [23], which confirms that at the time scale and length scale of the current simulations the twist diffusion can be considered instantaneous.

**Figure 1.**
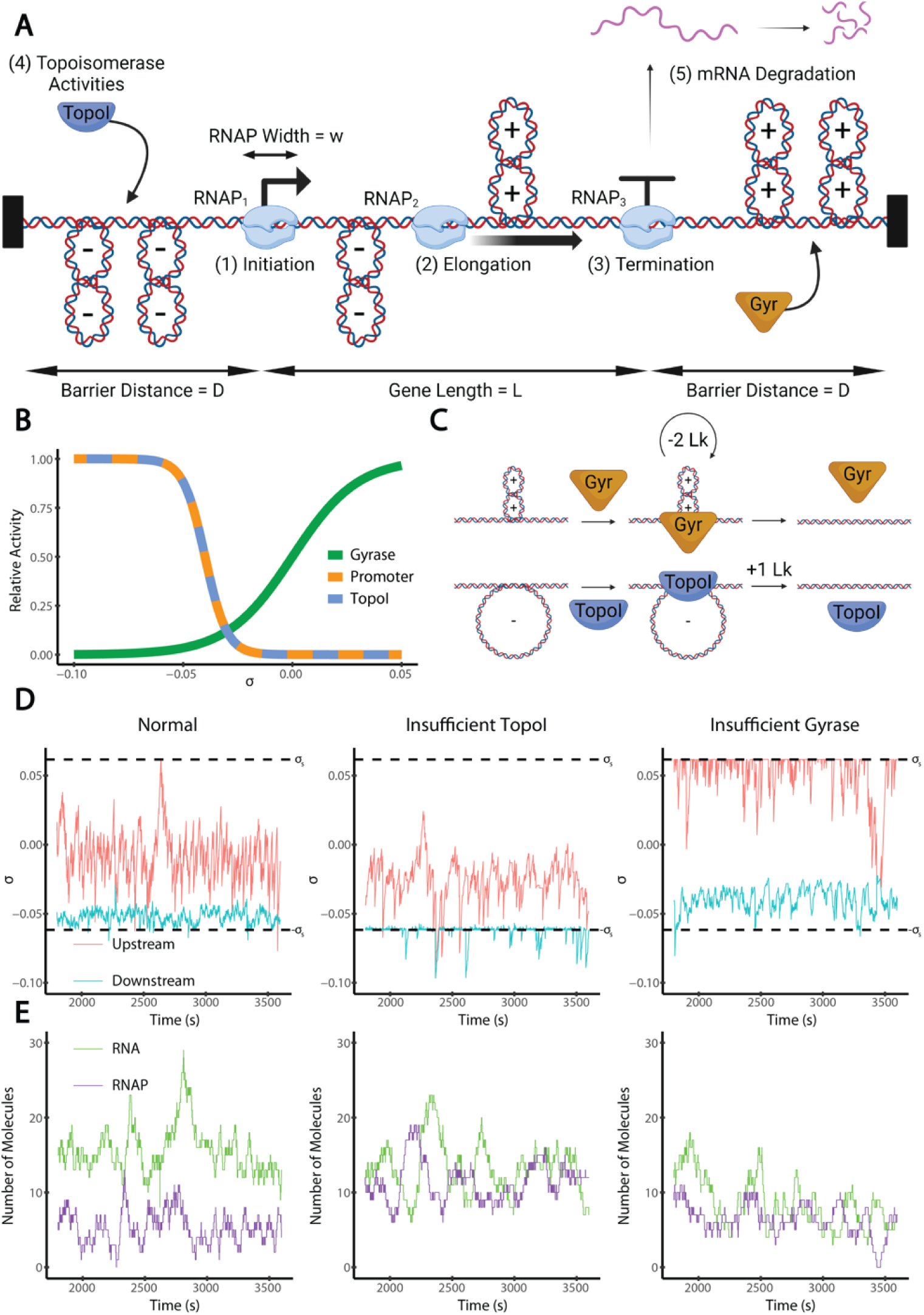
Model of Supercoiling-Coupled Transcription. (A) Model setup. For transcription, we consider the following fives aspects: (1) initiation, a single step process dependent on SC; (2) elongation, which introduces positive and negative supercoils downstream and upstream respectively; it occurs at a constant rate except when stalled under extreme supercoiling levels; (3) termination, which occurs instantaneously and produces a single RNA transcript; (4) topoisomerase activities, including gyrase and TopoI; and (5) RNA degradation, which occurs via a Poisson process. The gene length, L, is the distance between the transcription start site (TSS) and terminator; the barrier distance, D, is the sequence length between the TSS and upstream topological barrier and between the terminator and downstream topological barrier. (B) Modeled relative activity (normalized to the maximum possible rate) of gyrase binding (green), transcription initiation (orange), and TopoI binding (blue) at varying SC densities (σ). (C) Gyrase preferentially binds positively supercoiled DNA and decreases the linking number by 2 per catalytic cycle. Gyrase acts processively, performing 4 catalytic cycles per binding event on average. TopoI preferentially binds negatively supercoiled DNA and increases the linking number by 1 per binding event. (D) Example simulation time courses of supercoiling density (σ) upstream (blue) and downstream (red) of the gene under three different conditions, normal, insufficient Topo I activity, and insufficient gyrase activity. (E) Corresponding time courses of produced RNA copy number and number of bound RNAP molecules under the three conditions. The stalling SC thresholds, ±*σ_S_*, are shown as black dashed lines. Under the normal condition (left, gyrase activity 0.24 Lk/kb/s, TopoI activity 0.24 Lk/kb/s), transcription maintains continuous elongation with infrequent stalling. Under the insufficient TopoI activity condition (middle, gyrase activity 0.24 Lk/kb/s, TopoI activity 0.095 Lk/kb/s), transcription frequently stalls due to accumulated negative SC. Under the insufficient gyrase activity condition (right, gyrase activity 0.095 Lk/kb/s, TopoI activity 0.24 Lk/kb/s), transcription also stalls frequently due to accumulated positive SC. All examples are for strong genes (*k*_i_ = 0.2 s^-1^) with intermediately distant barriers (D = 10 kb).

On a stretch of DNA, SC is quantified by the difference between the linking number—the number of times the DNA strands wind around each other—of the DNA stretch and that of an equivalent length of relaxed B-form dsDNA. The linking number consolidates both the twist (the wrapping of DNA strands around their helical axis) and writhe (the wrapping of the DNA duplex along itself) into a single number; as both configurations present similar topological constraints to RNAP, and supercoiling density is only related to the change in the linking number, we do not consider the distinction between these two forms of SC.

We consider a list of DNA-bound RNAP molecules, (*RNAP*_1_, …, *RNAP*_k_), with *RNAP*_1_ being furthest upstream (or the newest RNAP molecule on the DNA). Each RNAP is considered to occupy a DNA stretch of length *w* = 30 bp to which other RNAP molecules cannot bind. For each *RNAP*_i_, we track the location of its center, *x*_i_, and the linking number of the DNA segment separating them from the nearest upstream RNAP_i-1_ or barrier, *Lk* ^(up)^, and the nearest downstream RNAP_i+1_ or barrier, *Lk* ^(down)^. From these linking numbers, we calculate a supercoiling density upstream (*σ* ^(up)^) and downstream (*σ*_i_ ^(down)^) of each RNAP molecule. The supercoiling density on a stretch of DNA is then defined by the formula

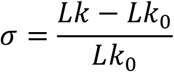

where *Lk* is the linking number of a DNA stretch and *Lk*_0_ is the linking number of the most relaxed state of the same length of DNA (10.5 bp^-1^). To manage the case where no RNAPs are bound, we also track *Lk*_dom_, the linking number of the domain. We initialize *Lk*_dom_ based on the initial supercoiling density *σ*_start_. In the absence of topoisomerases, *Lk*_dom_ is a constant value. Our simulation utilizes time steps of length *Δt* = 1/*v* where *v* = 25 bp/s is the RNAP velocity [3, 39].

### Initiation

We consider transcription initiation to occur in a single step facilitated by negative SC values. We assume that the concentration of RNAP is in great excess of DNA such that the concentration of the unbound RNAP pool stays constant. When initiation occurs, a new RNAP molecule is created with its center at the promoter position 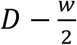, where *w* is the footprint of an RNAP molecule, so that the downstream end of the RNAP is at the transcription start site (TSS). This newly added RNAP is indexed as RNAP_1_, and the index of the existing RNAPs are increased by one (RNAP_1_ becomes RNAP_2,_ RNAP_2_ becomes RNAP_3_, etc.). Linking numbers are calculated and updated for the newly bound RNAP molecule and, if present, the adjacent downstream RNAP molecule. Initiation cannot occur if the promoter is occluded by an already-bound RNAP molecule, i.e. when RNAP(s) are bound and 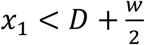. When the promoter is free, the initiation rate is dependent on the supercoiling density, *σ,* (Fig. 1B) according to the following sigmoidal function, where *k*_i_’ is the initiation rate; *k*_i_ is the maximal initiation rate (under infinitely negative SC); *σ*_i_ is the SC density at which *k*_i_’ is half of *k*_i_; and *β*_i_ is the crossover width, which characterizes the sensitivity of *k*_i_’ to changes in *σ*:

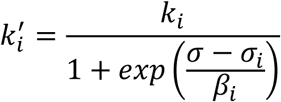

Based on *in vitro* data on transcription, we note that initiation reaches a maximum around *σ* = −0.05 and declines to a very low level around *σ* = −0.03 to *σ* = −0.01 [32]. We therefore use *σ*_i_ = −0.04 and *β*_i_ = 0.005 except where specified otherwise.

### Elongation

At each timestep, the position *x_i_* of each RNAP molecule (*RNAP_i_*) is increased by 1 bp, commensurate with a basal elongation speed of 25 bp/s in the absence of stalling. Stalling occurs for *RNAP_i_* at position *x*_i_ if *σ*_i_ ^(up)^ < −*σ*_s_ or *σ*_i_ ^(down)^ > *σ*_s_, where *σ*_s_ = 0.062 is the stalling threshold calculated from experimental observations [3, 17]. After each translocation, Supercoiling densities for the translocated and neighboring RNAP molecules are then recalculated based on their updated distances. The average elongation speed of each RNAP molecule is calculated by dividing the gene length by the time the RNAP molecule spends from initiation to termination. Thus, a frequently stalled RNAP molecule will have a slow elongation speed.

We note that some models have adopted a similar approach to model the stalling of elongating RNAP molecules [17], while others based stalling on the net torque experienced by RNAP, defined by a function of the difference between upstream and downstream superhelical densities [21, 40], and some use a combination of both approaches [23]. Experimentally, the upstream and downstream torques required to stall RNAP have been individually determined [3], but no similar data exists for torques exerted simultaneously upstream and downstream.

### Termination and RNA

Termination occurs deterministically and instantaneously when an RNAP molecule reaches the termination site, i.e., *x*_k_ = *L* + *D*. Each termination event increases the RNA copy number by one. Upon termination, *RNAP*_k_ is removed. If there are additional RNAP molecules bound to the gene, the linking number of the new downstream RNAP molecule after removal is updated to reflect the merged domains.

We assume that RNA molecules are degraded by a Poisson process, where each RNA molecule is assigned a lifetime drawn from an exponential distribution with expected lifetime *t*_RNA_, after which it is removed from the simulation. We use *t*_RNA_ = 120 s, on par with past experimental measurements [39].

We quantify transcriptional noise by calculating the Fano factor, calculated from the distribution of RNA copy numbers at each timepoint of the simulation by dividing the variance by the mean. A random birth-and-death Poisson transcription process has a Fano factor of one. Transcriptional bursting (bunched RNA production during “on” periods separated by inactive “off” periods) is characterized by a Fano factor greater than one.

### Topoisomerases

Topoisomerase activities are simulated stochastically. Gyrase and TopoI bind nonspecifically upstream and downstream of the gene. TopoI binds to the DNA at a rate determined by a decreasing sigmoidal function of SC defined by parameters *k*_T_, *σ*_T_, and *β*_T_, where *k*_T_ is the TopoI binding rate under optimal conditions, *σ*_T_ is the SC density where gyrase binds at half its maximum rate, and *β*_T_ is the crossover width of the SC dependency. The dependence of TopoI binding on SC density has not been fully determined experimentally. Therefore, except when noted otherwise, we take *σ*_T_ = *σ*_i_ = −0.04 and *β*_T_ = *β*_i_ = 0.005 on the grounds that both promoter opening and TopoI activity require DNA melting [8], as depicted in Fig. 1B. We did not model any direct interaction between TopoI and RNAP. Gyrase binds at a rate determined by an increasing sigmoid function of SC with analogous parameters *k*_G_, *σ*_G_, and *β*_G_. In line with available experimental data [37, 41], we take *σ*_G_ = 0 and *β*_G_ = 0.015, also shown in Fig. 1B.

When a TopoI molecule binds to a DNA stretch, it increases the linking number by one and unbinds instantaneously. This linking number is tracked both by any adjacent RNAP molecules and by the domain linking number (Lk_dom_).

When a gyrase molecule binds to a DNA stretch with a threshold of σ > *σ*_⍴_ = −0.11, it can perform multiple catalytic cycles before unbinding, as previously shown experimentally [36]. Gyrase processivity is simulated by drawing the number of catalytic cycles from a geometric distribution with mean ⍴_G_ = 4, where ⍴_G_ describes the average number of catalytic cycles per binding event [36]. We then cap the number of cycles such that gyrase never acts on DNA with σ < *σ*_⍴_ = −0.11; this value has been shown experimentally to be the lower bound on the SC level that gyrase can bind [37]. This procedure is equivalent to simulating successive catalytic cycles following the experimentally observed distribution of linking number changes introduced by gyrase [36]. Each catalytic cycle decreases the linking number by two, which is handled the same way as with TopoI.

When reporting “topoisomerase activities” for plotting purposes, we report 2*k_G_ρ_G_* as the gyrase activity and *k_T_* as the TopoI activity so that the rates are comparable in units of Lk/kb/s.

### Simulation

We simulated our model using a Gillespie algorithm [42] implemented in Python 3.9.18. Analysis and plotting were performed in R 4.3.1. Sections of simulation code were taken from Boulas et al. [17], and new sections were introduced to account for our treatment of initiation, topoisomerase activities, and RNA degradation. Parameter choices are indicated in Table 1.

**Table 1.**
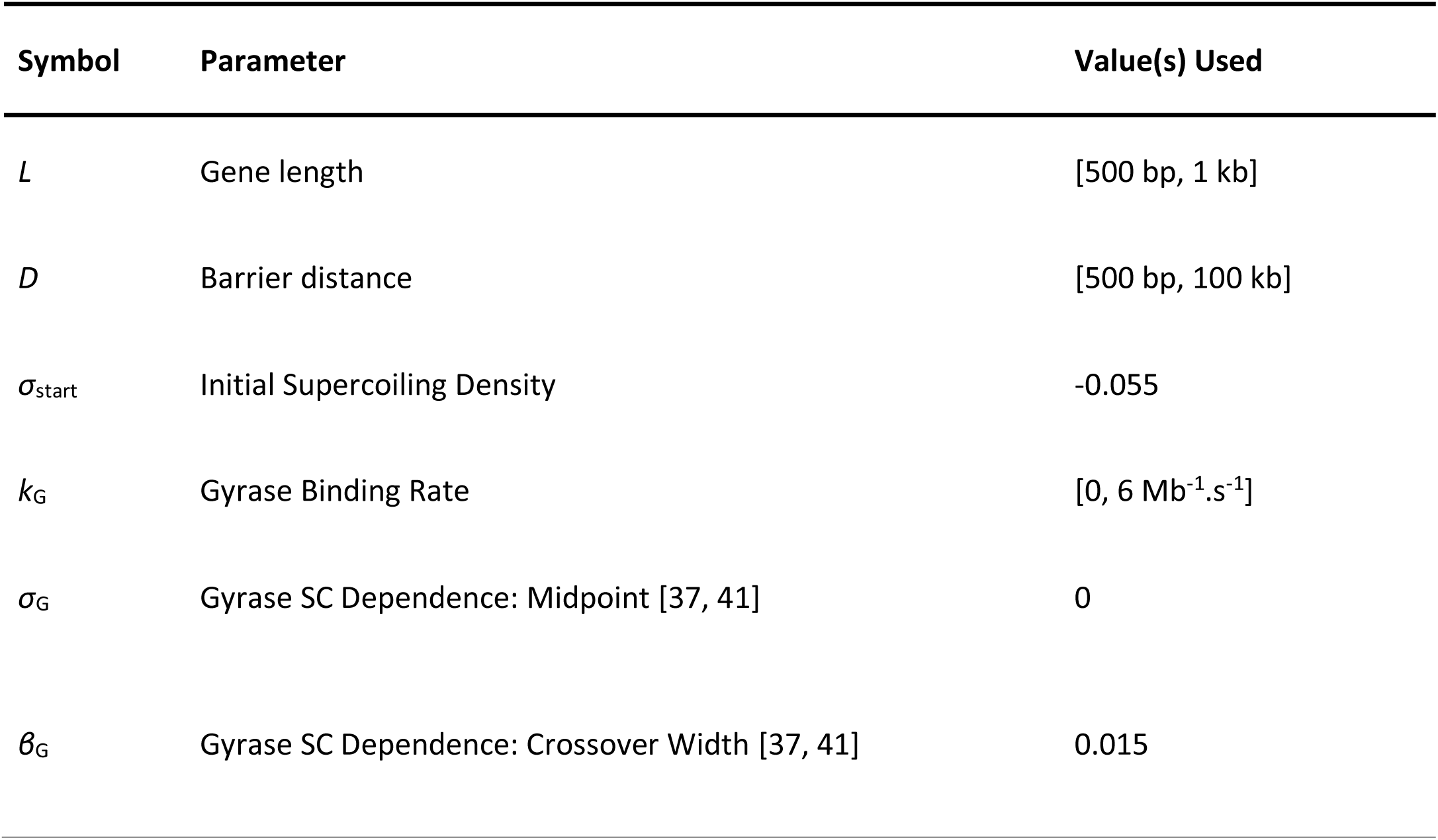
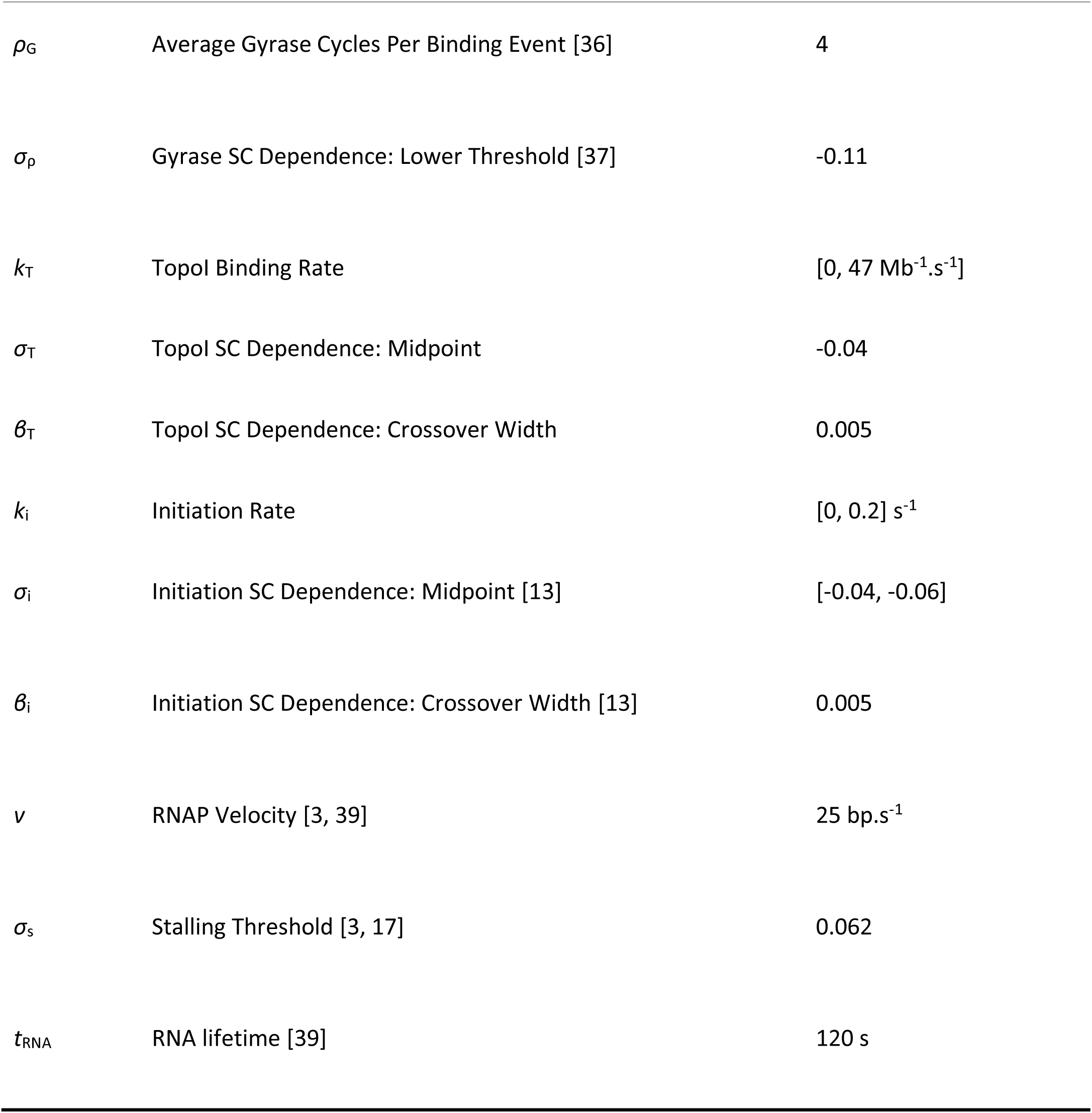
Parameter Definitions and Values.

| Symbol | Parameter | Value(s) Used |
| --- | --- | --- |
| $L$ | Gene length | [500 bp, 1 kb] |
| $D$ | Barrier distance | [500 bp, 100 kb] |
| $\sigma_{\text{start}}$ | Initial Supercoiling Density | -0.055 |
| $k_G$ | Gyrase Binding Rate | [0, 6 Mb<sup>-1</sup>.s<sup>-1</sup>] |
| $\sigma_G$ | Gyrase SC Dependence: Midpoint [37, 41] | 0 |
| $\beta_G$ | Gyrase SC Dependence: Crossover Width [37, 41] | 0.015 |
| $\rho_G$ | Average Gyrase Cycles Per Binding Event [36] | 4 |
| $\sigma_p$ | Gyrase SC Dependence: Lower Threshold [37] | -0.11 |
| $k_T$ | Topol Binding Rate | [0, 47 Mb<sup>-1</sup>.s<sup>-1</sup>] |
| $\sigma_T$ | Topol SC Dependence: Midpoint | -0.04 |
| $\beta_T$ | Topol SC Dependence: Crossover Width | 0.005 |
| $k_i$ | Initiation Rate | [0, 0.2] s<sup>-1</sup> |
| $\sigma_i$ | Initiation SC Dependence: Midpoint [13] | [-0.04, -0.06] |
| $\beta_i$ | Initiation SC Dependence: Crossover Width [13] | 0.005 |
| $v$ | RNAP Velocity [3, 39] | 25 bp.s<sup>-1</sup> |
| $\sigma_s$ | Stalling Threshold [3, 17] | 0.062 |
| $t_{RNA}$ | RNA lifetime [39] | 120 s |

### Transcription of a freely rotating plasmid without topoisomerases

We simulated the case of a gene transcribed on a circular plasmid without topoisomerases (as in a usual *in-vitro* setup), where positive and negative SC introduced by elongation diffuse around the plasmid and annihilate one another. This leads to a constant global SC level. We simulated short genes (*L* = 500 bp) with a strong promoter (*k*_i_ = 0.4 s^-1^) that opens at different SC densities (*σ*_i_ = −0.06, −0.05, and −0.04). Since our simulation scheme was designed for linear DNAs, we approximated a circular plasmid by using extremely distant barriers (*D* = 10^9^ bp), resulting in negligible SC accumulation. For each condition, we ran 21 simulations, varying the SC density from *σ*_start_ = −0.1 to *σ*_start_ = 0 in steps of 0.005. Simulations were run for 10^6^ timesteps (11.11 hours simulated time), using approximately 1.7 CPU-minutes each. To calculate the transcription rate (number of RNA/time), we divided the number of termination events (i.e., the number of produced RNA) by the total time of each simulation for individual plasmid topoisomers. To compare the simulations with transcription levels from a distribution of topoisomers, we assumed a normal distribution of linking numbers with a standard deviation of 0.004, a typical value for plasmids extracted from cells and measured by agarose-chloroquine gels [13]. We then calculated the transcription rates for distributions of plasmid topoisomers with mean SC densities ranging from −0.1 to 0.

### Transcriptional regulation of a topologically isolated gene

We simulated transcription of a gene on linear DNA flanked by topological barriers in the presence of topoisomerases, mimicking a topological domain *in vivo*. In all simulations, we used a gene length *L* = 1 kb. We varied four variables between these simulations: promoter strength (*k*_i_), barrier distance (*D*), TopoI binding rate (*k*_T_), and gyrase binding rate (*k*_G_). Promoter strengths were chosen based on the expected number of RNAP molecules on the gene assuming conditions permissive of initiation and elongation to simulate different transcription regimes: a weak promoter (k_i_ = 0.008 s^-1^, average RNAP molecules per gene = 0.32) simulates intermittent transcription; a moderate promoter (k_i_ = 0.05 s^-1^, avg. RNAP molecules per gene = 2) simulates mostly continuous transcription with occasional breaks; and a strong promoter (k_i_ = 0.2 s^-1^, avg. RNAP molecules per gene = 8) simulates continuous transcription. As a control, we also consider the case of no transcription. We consider barrier distances of *D* = 1 kb (close barriers), 10 kb (intermediate barriers), and 100 kb (far barriers).

For each barrier distance, we calculate the steady-state binding rates of both TopoI and gyrase such that TopoI removes the upstream negative SC and gyrase removes the downstream positive SC at the same rate SC is introduced by elongation, assuming TopoI and gyrase bind at their maximum rates. We term these values *k*_T_* and *k*_G_*, and we use these values to guide our parameter choices, as we would expect different behaviors between *k*_T_ < *k*_T_* and *k*_T_ > *k*_T_*, and likewise between *k*_G_ < *k*_G_* and *k*_G_ > *k*_G_*. We simulate each combination of *k*_T_ and *k*_G_ achieved by varying *k*_T_ from 0 to 2*k*_T_* in increments of 0.1*k*_T_* and *k*_G_ from 0 to 2*k*_G_* in increments of 0.1*k*_G_*. Note that *k*_T_* and *k*_G_* are inversely proportional to the barrier distance *D* (since we assume non-specific binding of topoisomerases). We also run a second set of simulations at 1 kb and 100 kb barrier distances with the same *k_T_* and *k*_G_ ranges used for the 10 kb barrier distance to enable comparisons between barrier distances at the same nonspecific topoisomerase activities.

We therefore generated 2420 total simulations, examining 11 TopoI conditions, 11 gyrase conditions, 4 promoter conditions, and 5 barrier conditions (2 topoisomerase ranges for close and far barrier distances, 1 topoisomerase range for intermediate barrier distances). Simulations were run for 2 × 10^6^ timesteps (22.22 hours simulated time), utilizing approximately 4 CPU-minutes per simulation. In most cases, we disregard the first 45,000 timesteps (30 minutes) to focus on the steady-state behavior of transcription (in cases with 100 kb barrier distances, we had to remove 60 minutes of data, as the simulations took longer to equilibrate). To isolate the impact of topological variables on transcription from the strength of the promoter, we calculate a normalized transcription rate by dividing the observed average transcription rate by the maximal promoter initiation rate (*k*_i_). We also calculate a “free promoter fraction”, representative of the proportion of time steps during which the promoter is not blocked from binding by an RNAP. Low free promoter fraction indicates that transcription is repressed by slow elongation, as RNAP does not clear the promoter in a timely manner. This value is calculated by dividing the observed transcription rate by the mean value of the effective initiation rate given the promoter SC density (*k*_i_’). In our model, all RNAPs that initiate eventually produce an RNA transcript and initiation is determined exclusively by the promoter SC density and the occupancy of the promoter by RNAP, so this value isolates the effect of promoter occupancy and characterizes the fraction of time during which the promoter is free for RNAP binding.

## Results

### Simulation of *in vitro* transcription recapitulates optimal expression at physiological negative superhelical levels

We first compared the predictions of our model to *in vitro* transcription data obtained using circular plasmids of different superhelical densities incubated with RNAP. These data have been a major source for identifying the SC-sensitivity of bacterial promoters [30–32]. Because supercoils generated by transcription can diffuse and annihilate each other in a circular freely rotating plasmid, the SC level can be approximated as constant in such templates. Previous experiments have shown that a gene typically transcribes maximally at an optimal negative SC level and exhibits decreased transcription when the negative SC level is too low or too high (S1 Fig). However, no stochastic model has reproduced this non-monotonic behavior, a basic yet crucial observation of transcription in response to negative SC.

In our simulations, transcription ceased completely at very negative SC densities (*σ* < *σ_s_*) due to RNAP stalling (Fig. 2A, left side of the red vertical line), and decreased continuously when the SC density increases above this threshold (Fig 2A, right side of the red vertical line), due to a reduced initiation rate. This transcriptional response to the SC density of the plasmid holds true for promoters with different initiation dependencies on SC (*σ*_i_, Methods) (Fig. 2A, compare data points of different colors).

**Figure 2.**
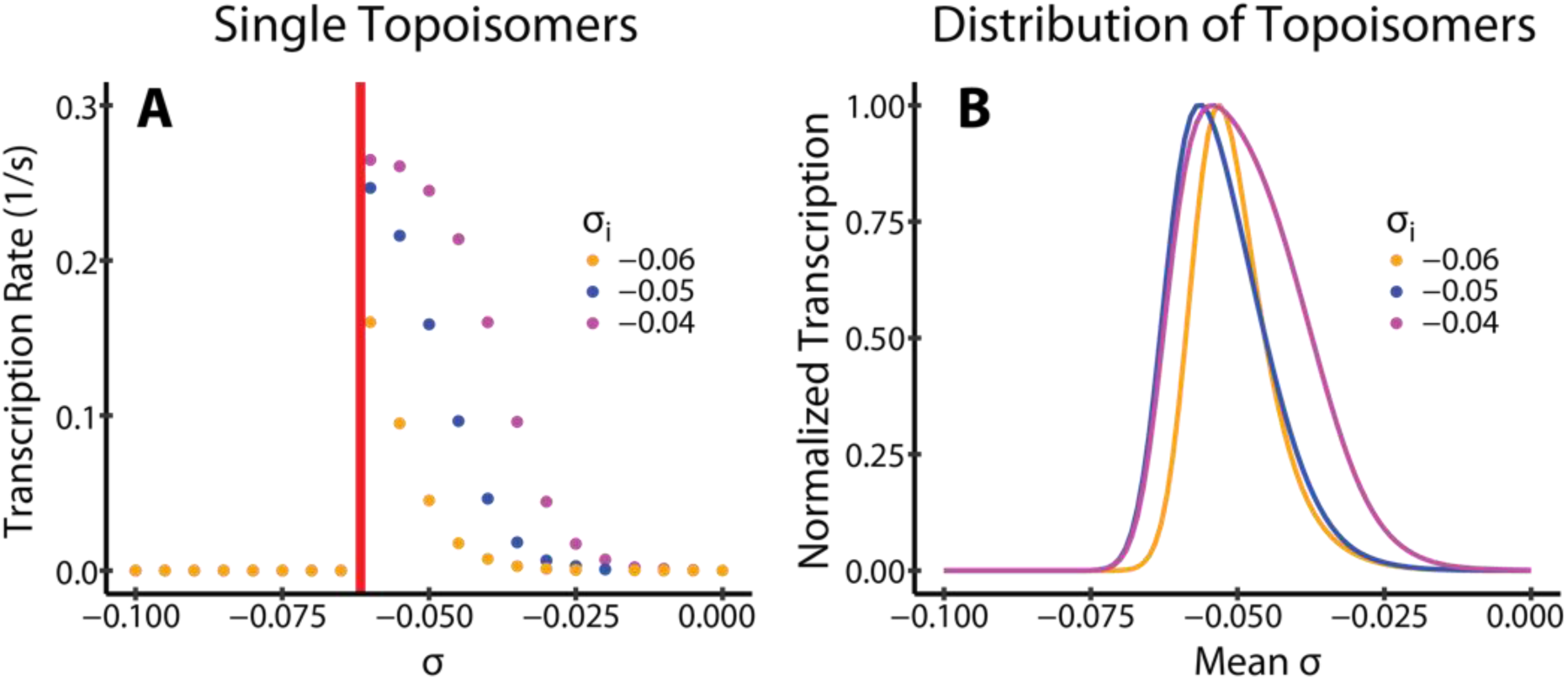
Transcription on Freely Rotating Plasmids. We simulate transcription on freely rotating plasmids in the absence of topoisomerases, mimicking *in vitro* transcription assays. (A) Transcription rates of three promoters that open at different *σ*_i_, simulated on plasmids with different SC densities, σ, but with only a single topoisomer. The RNAP stalling threshold *σ_s_*is indicated in red. (B) Transcription rates of the same three promoters simulated on plasmid samples with normally distributed SC densities with multiple topoisomers, which mimics typical topoisomer [30–32] distributions of plasmids extracted from cells.

To further compare with experiments performed on plasmid samples of different mean SC densities and with a distribution of topoisomers (with different linking numbers, as commonly extracted from cells), we computed the total expression level from the weighted averages of different templates (each corresponding to one plasmid topoisomer), and calculated the effective transcription rate of samples with different mean SC densities. Fig. 2B shows the resulting continuous SC-dependency where transcription is maximized by a moderately negative SC density, and decreased in either direction from this optimum, recapitulating previous experimental observations [30–32]. This behavior is observed regardless of the choice of *σ*_i_, and is therefore valid for different promoters. These simulation results suggest that our model can faithfully recapitulate the non-monotonic dependence of transcription on negative SC levels. The very sharp decrease observed at strongly negative superhelical levels in our simulations (compared to the datapoints shown in S1 Fig) suggests that in *in vitro* transcription systems, the frictional drag experienced by RNAP might be weaker due to the absence of translation and a less crowded environment compared to the model assumption.

### Regulatory effects of topoisomerases’ activities depend on promoter strength

Next, we simulated transcription on linear DNAs with topological barriers in the presence of topoisomerase activities. This situation mimics typical transcription occurring in bacterial chromosomal topological domains. We simulated topological barriers at distances of 10 kb flanking a typically sized gene of 1 kb, leading to a 21 kb topological domain, in line with experimentally reported sizes in *E. coli* [43]. TopoI and gyrase molecules can bind upstream or downstream of elongating RNAPs depending on their binding rates and the local SC densities (see Methods). In the absence of transcription, the relative activities of the two topoisomerases maintain a domain-wide (homeostatic) SC level of about σ = −0.06 to −0.02, consistent with that typically measured in bacterial cells (S2 Fig, A). Meanwhile, transcription substantially alters the spatiotemporal distribution of SC in the domain (S2 Fig, C) while maintaining a similar average domain-wide SC density (S2 Fig, B).

In our simulations, we define the transcription rate as the number of RNA molecules generated per unit time, normalized to the maximal transcription rate that promoter can achieve (Fig. 3, row A). Note as we do not consider premature termination in our simulations, all RNAP molecules that initiate will eventually produce a transcript. Therefore, the transcription rate is equivalent to the transcription initiation rate and should be independent of the elongation speed (Fig. 3, row B). However, an RNAP molecule may stay at the promoter and occlude the binding of the next RNAP molecule to the promoter, if it is stalled at the promoter due to an unfavorable SC level or blocked from moving past the promoter due to piled, stalled RNAP molecules in the gene body. Therefore, a slow elongation speed, which reflects stalling during initiation and/or elongation, could impact the transcription rate through the availability of the promoter to the next RNAP molecule. To illustrate this point, we calculated the free promoter fraction, which is the proportion of time steps during which the promoter is free for RNAP binding (Fig. 3, row C). As such, the free promoter fraction is a direct indicator of the impact of RNAP’s stalling on the transcription rate, with lower free promoter fractions indicating transcription repression caused by stalled RNAP.

**Figure 3.**
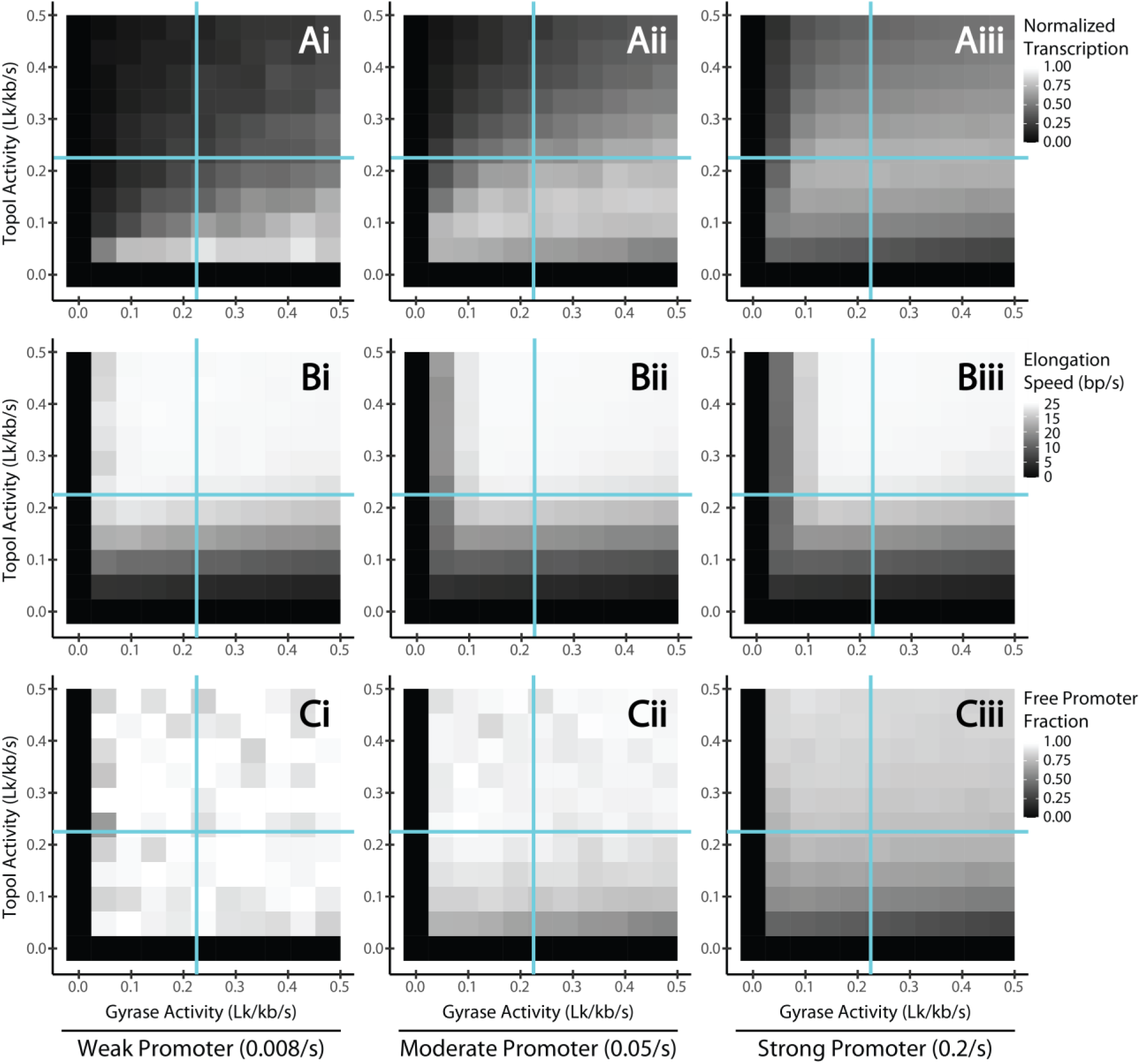
Regulatory Effect of Topoisomerases Activities on Transcription. Mean normalized transcription rate (each value is *k*_obs_/*k*_i_, i.e. normalized by the initiation rate of the promoter in the absence of supercoiling constraints, row A), mean elongation speed (bp/s, row B), and the free promoter fraction (the proportion of timesteps during which the promoter is not sterically occluded by a bound RNAP molecule, row C) are plotted as two-dimensional heatmaps at different combinations of Topo I (y-axis) and gyrase (x-axis) activities. We simulated 1 kb genes flanked by 10 kb barriers with three promoter strengths, weak, moderate and strong (columns i, ii, and iii respectively). The intersections of the two cyan lines denote the steady-state topoisomerase activities needed to cancel SC generated by elongation.

From these simulations, we observe that TopoI and gyrase impact promoters of varied strengths in a qualitatively different manner, with each promoter reaching its maximal transcription rate at a different combination of TopoI and gyrase activities (Fig. 3, row A, compare the positions of the brightest squares in heatmaps for weak, moderate and strong promoters). Specifically, for a weak promoter, high gyrase activity (leading to more negative SC densities) increases the transcription rate, while a high TopoI activity (leading to less negative SC density) inhibits the transcription rate (light grey and white squares in Fig. 3Ai). Interestingly, the highest transcription rate of the weak promoter occurred at the slowest elongation speed (dark grey squares in Fig. 3Bi). In other words, slow elongation speed, which signals frequent RNAP stalling, does not appear to reduce the overall transcription rate for the weak promoter. However, as the free promoter fractions across different topoisomerase conditions remain high (light grey and white squares in Fig. 3Ci), it indicates that the weak promoter is limited by transcription initiation but not RNAP stalling. The promoter is maximally transcribed under topoisomerase conditions that result in the most negative SC (low TopoI and high gyrase activities, white squares in Fig. 3Ai), which facilitate promoter opening and elongation to increase the transcription rate. Under this condition, RNAP stalling due to insufficient topoisomerases’ activities still occurs, however it does not substantially affect the transcription rate because the initiation rate is limiting.

A strong promoter (Fig. 3, column iii) displays a markedly different behavior. Maximal transcription rate was reached at a moderate activity of TopoI and moderate to high activities of gyrase (light grey squares in Fig. 3Aiii), which corresponds to relatively high elongation speeds, i.e., low levels of RNAP stalling (light grey squares in Fig. 3Biii) and moderate free promoter fraction (grey squares in Fig. 3Ciii). Too high or too low TopoI activities reduced the overall transcription rate regardless of gyrase activity. At high TopoI activities, reducing gyrase activity represses transcription (top dark grey squares in Fig. 3Aiii), which corresponds to substantially decreased elongation speeds, i.e., frequently stalled RNAP (top white to dark grey squares in Fig. 3Biii), but not dramatically lowered free promoter fractions (top light grey squares in Fig. 3Ciii). These observations show that in this case, the weakly negative SC level at the promoter reduces initiation, while RNAP stalling along the gene has no consequence for the transcription rate. Low TopoI activity and, to a lesser degree, low gyrase activity, decrease both the elongation speed and free promoter fraction (dark grey and black squares in Fig. 3Biii, Ciii), indicating insufficient topoisomerase activities to remove SC introduced by transcription, which leads to stalled RNAP molecules both at the promoter and during elongation. Thus, in moderate to low TopoI activities, a strong promoter is more sensitive to RNAP-stalling-mediated transcription repression than a weak promoter due to its more frequent RNAP binding at the promoter.

A moderate promoter (Fig. 3, column ii) displays characteristics of both strong and weak genes. In conditions of low TopoI activities, it behaves similarly to the strong gene, where low TopoI levels result in repressed transcription (Fig. 3Aii) as elongation is slowed (Fig. 3Bii) and the free promoter fraction decreases (Fig. 3Cii). At high TopoI levels, the moderate promoter behaves similarly to the weak promoter case, with a consistently high free promoter fraction (Fig. 3Cii) and high elongation rate at high gyrase and low TopoI activities.

Taken together, our simulations show that the transcription rate can be regulated by combined Topo I and Gyrase activities through the availability of the promoter. A high free promoter fraction as that in the case of the weak promoter signals that the promoter SC level is limiting for RNAP to initiate transcription, while a low free promoter fraction as that in the case of the strong promoter suggests that RNAP stalling is limiting for making the promoter available for the next RNAP molecule. Specifically, in conditions where the SC-dependent initiation rate, *k*_i_’, is low, the free promoter fraction is high, and transcription is regulated primarily by the promoter SC level for initiation. Therefore, transcription is enhanced by increasing gyrase activity and decreasing TopoI activity to increase negative SC for promoter opening. As *k*_i_’ increases, the increased frequency of RNAP binding means that the free promoter fraction becomes sensitive to RNAP stalling. Thus, the transcription rate becomes highly dependent on the presence of adequate levels of both TopoI and gyrase to keep up with the SC introduced by elongation to avoid stalling. With high *k*_i_’, the favorability of the promoter SC level for initiation becomes relevant only when comparing multiple conditions in which TopoI and gyrase are sufficient for avoiding stalling. In general, fast initiation leads to transcription being dependent on RNAP stalling, while slow initiation leads to transcription being dependent on the promoter SC level.

### Regulation by topological domain size

We next investigated the dependency of transcription on topological domain size. Specifically, we reduced the barrier distance to investigate the effects of imposing a topological constraint, similar to that in a DNA loop formation due to the binding of proteins [46–48]. We kept the same gene length (1 kb) and promoter strengths but reduced the barrier distances from 10 kb to 1 kb (corresponding to a total domain size of 3 kb instead of 21 kb). We then calculated the fold-change in the transcription rate resulting from reducing the barrier distance (Fig 4A). We observed that for a strong promoter, a smaller domain size repressed transcription (blue squares in Fig. 4Aiii), while for weak and moderate promoters, a smaller domain size has varied effect on transcription (red and blue squares in Fig. 4Ai, Aii): it represses transcription in the region of low TopoI and high gyrase activities, where a higher level of negative SC in the larger domain facilitates initiation, and activates transcription in the region of high TopoI and low gyrase activities, where a less negative SC level in the larger domain inhibits transcription initiation. We note that these effects are quite similar to those observed earlier after reducing both TopoI and gyrase activity (Fig 3, row A). This similarity was expected as in our model, topoisomerases act nonspecifically, meaning that proportionally fewer binding events of topoisomerases occur on the smaller domain. This situation mimics that in cells where chromosomal domain forms in the presence of the same cellular pool of topoisomerases. We confirmed that the observed change in transcription is indeed primarily due to the decreased effective topoisomerase binding: when we increased the nonspecific topoisomerase binding rates tenfold (thus keeping their effective binding approximately constant), we observed that the effect of the small domain size is largely eliminated (Fig. 4B).

**Figure 4.**
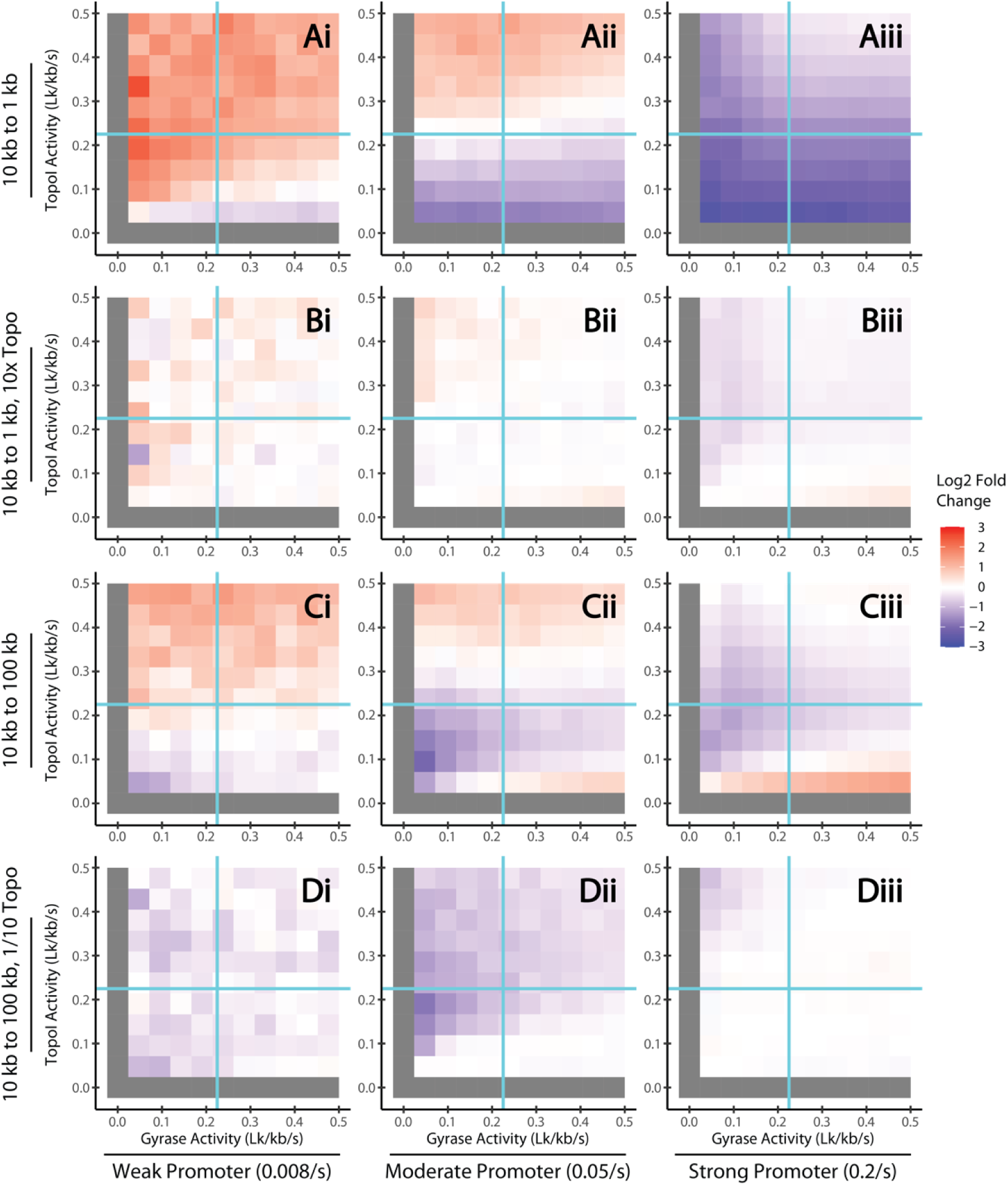
Regulatory Effect of Topological Domain Size. The log2 fold changes in the transcription rate between two simulations (red: higher; blue: lower) for weak (column i), moderate (column ii), and strong (column iii) promoters were plotted across a range of TopoI and gyrase activities (Lk/kb/s). (A) A 10 kb barrier distance is decreased to 1 kb. (B) The domain sizes are the same as that in (A) except that the nonspecific topoisomerase activities are increased tenfold as the barrier distance is decreased. (C) A 10 kb barrier distance is increased to 100 kb. (D) The domain sizes are the same as that in (C) except that the nonspecific topoisomerase activities are decreased tenfold as the barrier distance is increased. In all cases, the displayed TopoI and gyrase activities correspond to those of the initial barrier distance and interaction of cyan lines denotes the topoisomerase activities calculated to eliminate the supercoils introduced by elongation for the initial barrier distance. For simulations with 100 kb barriers the first hour (rather than 30 minutes) of simulation time was disregarded due to longer transient states.

Next, we increased the barrier distance from 10 kb to 100 kb and observed a complex response of the transcription level with a non-monotonic dependence on both gyrase and TopoI activities (Fig 4C). Because this scenario mimics the opening of a chromosomal domain and the dissipation of supercoiling constraints in a larger domain, a gene with distant barriers therefore displays a transcription rate that is only weakly dependent on topoisomerase activities (S3 Fig, row A), compared to a gene surrounded by intermediate barriers (Fig. 3A). A large domain allows sufficient topoisomerase activities to avoid RNAP stalling (S3 Fig, row B) and leads to a high free promoter fraction (S3 Fig, row C) under all tested promoter strengths and topoisomerase binding conditions. Therefore, the fold-change in response to an increased barrier distance closely resembles the inverse of the transcription rate with intermediate barriers (Fig 3A). The exception to this trend is for the moderate and strong genes, at very low TopoI and high gyrase activities, where increasing the barrier distance activates transcription due to a very negative promoter SC density (S3 Fig, row A), which is repressive for the same moderate and strong genes in the case of intermediate barriers. As with the barrier reduction, we confirm that this result is mediated by the effective topoisomerase binding rates across the domain, as decreasing the topoisomerase binding rates tenfold removes these trends (Fig 4D).

Taken together, these observations suggest that the size of a topological domain imposes an additional layer of transcription regulation, primarily through the availability of topoisomerases in the domain. The formation of a smaller topological domain reduces the number of topoisomerase binding events, altering the local SC level relevant to the genes within the domain. Strong genes in smaller domains are hence repressed by the inability of topoisomerases to keep up with the SC introduced by elongation. Meanwhile, weak to moderate genes respond differentially depending on the combination of the two topoisomerase activities. On large domains, transcription is favored by low TopoI and high gyrase levels, as the effective topoisomerase activities increase, resulting in a high free promoter fraction. The interaction between promoter strength, topoisomerase binding rates, and topological context must therefore be considered to quantitatively understand a gene’s regulation.

### SC modulation of the off-state leads to transcriptional bursting

Transcriptional bursting is the phenomenon of a gene possessing multiple states in which transcription occurs at different rates (typically on or off) [49]. The bursting phenomenon results in transcriptional noise greater than that of a random birth-and-death Poisson process, typically inducing variable gene expression in an isogenic cell population, which enhances its resilience to environmental uncertainty [28]. Since SC modulates transcriptional initiation in our model, we wished to understand whether it could serve as a generator of transcriptional bursting in the absence of additional factors such as stochastic binding of regulators or other proteins affecting the promoter accessibility. We therefore analyzed the simulations to characterize the conditions that result in transcriptional bursting and identify underlying mechanisms.

The level of noise can be quantified using the normalized variance of expression (variance divided by the square of the mean), but this parameter decreases with the expression level (S4 Fig.), making it difficult to compare simulations involving different promoter strengths. Rather, we computed Fano factors (variance divided by the mean) across all promoter strengths, topoisomerase binding rates and barrier distances. A Fano factor of one indicates Poissonian expression (i.e., constant initiation rate), whereas values greater than one indicate the presence of transcriptional bursting. We observed Fano factor values ranging from 0.7 to 4.8, with bursty transcription appearing in diverse promoter and topological conditions (Fig. 5A). The datapoints separate horizontally along the axis of mean RNA copy number into three clouds corresponding to simulations with weak, moderate and strong promoters, respectively. For most conditions, the distance to barriers has little impact on transcription burstiness—the clouds of red, green and blue datapoints for close, intermediate and far distances respectively are mostly close to each other. However, promoter strength appears to play a role. Weak promoters’ transcription is close to Poissonian (Fano factors close to 1, squares, Fig. 5A), with moderate promoters exhibiting consistently higher Fano factors (about 1-2, triangles, Fig. 5A) indicative of moderate burstiness. For strong promoters (circles, Fig. 5A), we observe a strong increase of Fano factors specifically for a group of simulations with high barrier distance (blue dots, D = 100 kb), high TopoI activity, and low gyrase activity (*i.e.*, in a topologically very relaxed domain, S5 Fig.). Interestingly, for all promoters, and especially for strong promoters, we observed a large number of simulations exhibited Fano factors below 1*, i.e.,* sub-Poissonian, suggesting that transcription becomes less random because initiation events are more regularly spaced by the time RNAP molecules take to clear the promoter.

**Figure 5.**
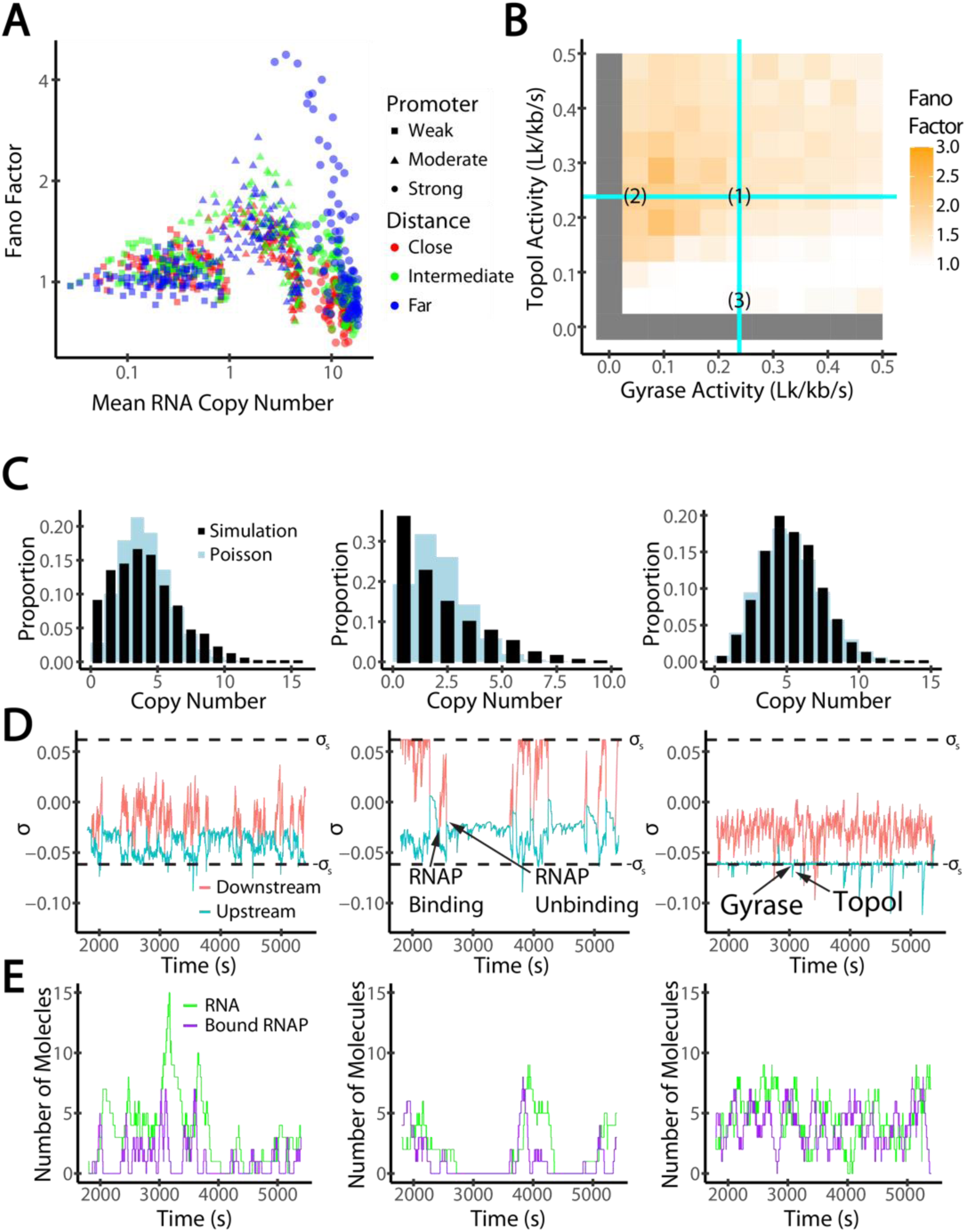
Mechanism of Transcriptional Bursting. (A) Fano factors and mean RNA copy numbers for simulations run squares, *k*_i_ = 0.008 s^-1^, moderate: triangles, *k*_i_ = 0.05 s^-1^, strong: circles, *k*_i_ = 0.2 s^-1^), barrier distances (close: red, *D* = 1 kb, intermediate: green, *D* = 10 kb, far: blue, *D* = 100 kb), and topoisomerase binding rates (one point per simulation). Data are shown on log-log scale. (B) Fano Factors for a moderate gene (*k*_i_ = 0.05 s^-1^) flanked by intermediate barriers (*D* = 10 kb). Squares correspond to individual simulations with varied topoisomerase activities. Cyan lines denote the steady-state topoisomerase activities calculated to eliminate the supercoils generated by elongation. Scenarios (1), (2), and (3) are analyzed in the remainder of the figure. (C) Distributions of RNA copy numbers (black) throughout one simulation compared with the Poisson distribution with the same mean (blue). (D) Time course of the SC density upstream (blue) and downstream (red) of the gene. The stalling thresholds, ±*σ_S_*, are indicated with black dashed lines. Features relevant to transcriptional bursting are indicated, including an RNAP molecule binding to the gene and an RNAP molecule unbinding after completing transcription (middle) and gyrase and Topo I binding (right). Note that RNAP unbinding leads to the merge of the upstream (red) and downstream (cyan) SC domains. (E) The time course of copy numbers of RNA (green) and bound RNAP (purple) molecules.

To further explore transcription bursting dynamics, we consider in-depth the case of a moderate promoter (k_i_ = 0.05 s^-1^) with intermediate barriers (D = 10 kb). This promoter strength and domain size (green triangles in Fig. 5A) are representative of many bacterial genes and exhibit weakly bursty transcription independent of barrier distance (S4 Fig), matching experimental observations of Fano factors typically between 1 and 3 for native genes [28]. In Fig. 5B we plotted Fano factors of these simulations at different combinations of TopoI and gyrase activities and chose three scenarios for detailed analyses. In scenario 1, the Fano Factor is 1.6 and the steady-state topoisomerase binding rates exactly cancel the SC introduced by elongation; In scenario 2, the Fano Factor is at 2.0 and the gyrase activity is lower than that in Scenario 1; In scenario 3, the Fano Factor is 0.88 and the TopoI activity is lower than that in Scenario 1. For each scenario, we consider the distribution of the RNA copy numbers (Fig 5C), the time course of the upstream and downstream SC density (Fig 5D), and the time course of the RNA copy number and number of bound RNAP molecules (Fig 5E).

Scenarios 1 and especially 2 describe a mechanism for transcriptional bursting in which the upstream negative SC induced by elongation facilitates the binding of additional RNAP molecules at the promoter, hence facilitating more frequent initiation events. In scenario 2, very clear “on-off” transitions are observed in accordance with a higher Fano factor at 2: the very low gyrase activity resulted in a relatively high SC density at the promoter that mostly, but not completely, suppresses initiation (the “off” state) when the gene is not bound by any RNAP molecules, whereas a single transcription event turns the gene into the “on” state by reducing the promoter SC level (Fig 5D-E). Whenever all RNAP molecules stochastically unbind, the positive SC accumulated downstream diffuses onto the promoter, suppressing initiation and returning the gene to the off state as the existing RNA fully decay. Scenario 1 demonstrates on- and off-state by the same mechanism, but the off-state is shorter as the relatively high gyrase activity results in an steady-state SC level more favorable for initiation before previous RNA molecules are degraded in the off-state (Fano factor of 1.60). This mechanism implies a cooperative mode of multiple RNAP molecules as predicated previously and is interesting for its alignment with the experimental observation that transcriptional bursting is related to low gyrase expression [28]. Scenario 2 displays a greater Fano factor than scenario 1 (2.0 vs 1.6) because the greater steady-state SC level in the absence of bound RNAP molecules results in less frequent RNAP binding and more prolonged off-state.

In scenario 3, we do not observe transcriptional bursting; in fact, we observe a sub-Poisson Fano factor (0.88). The SC level is strongly negative, almost always sustaining efficient initiation, hence a high transcription level, but so much so that elongation frequently stalled due to insufficient TopoI activity. Therefore, several RNAPs are almost always bound to the gene, and the promoter never reaches an “off” state. Because these abundant RNAPs frequently block the promoter from binding additional RNAPs, initiation events are more evenly spaced than in a Poisson distribution, explaining the low Fano factor.

We conclude that transcriptional bursting is dependent on a steady-state SC level in the “off” state which represses, although does not eliminate, initiation, while negative SC induced by elongation turn the promoter into a transitory “on” state (*i.e.*, transcription is self-activating).

### Transcriptomics data support promoter strength-dependent regulation by topoisomerases

Our modeling effort so far demonstrated that combinations of topoisomerase activities could lead to differential effects on transcription, independent of promoter-specific regulators. To verify whether these impact hold for broad sets of genes *in vivo*, we compiled previously published transcriptomic datasets [9–17, 33] obtained in evolutionarily distant bacterial species (S1 Table) where cells were treated with a gyrase inhibitor (fluoroquinolones or coumarins) or TopoI inhibitor (seconeolitsine).

Because the datasets are heterogeneous in the type of species, culture conditions, drug dosages, treatment time, and experimental methods (in particular microarrays vs RNA-Seq), we first grouped genes into expression quartiles to improve readability. In Fig. 6A, we illustrated typical relations between gene expression strength and transcriptional response to gyrase inhibition, for *Escherichia coli* and *Mycoplasma pneumoniae*. In both cases, we clearly observed that genes of higher expression associated with transcriptional repression when gyrase is inhibited, albeit the effect has a very different magnitude in the two species (the absolute expression ratio between the first and last quartiles is around 4-fold for *E. coli* vs around 30-fold for *M. pneumoniae*). In contrast, genes of the weakest quartile showed activation. Note that in these analyses, the expression strengths are normalized to the total transcriptional output, so it cannot be concluded that weak genes are upregulated, but rather that their share in total mRNA numbers has increased after the treatment.

**Figure 6.**
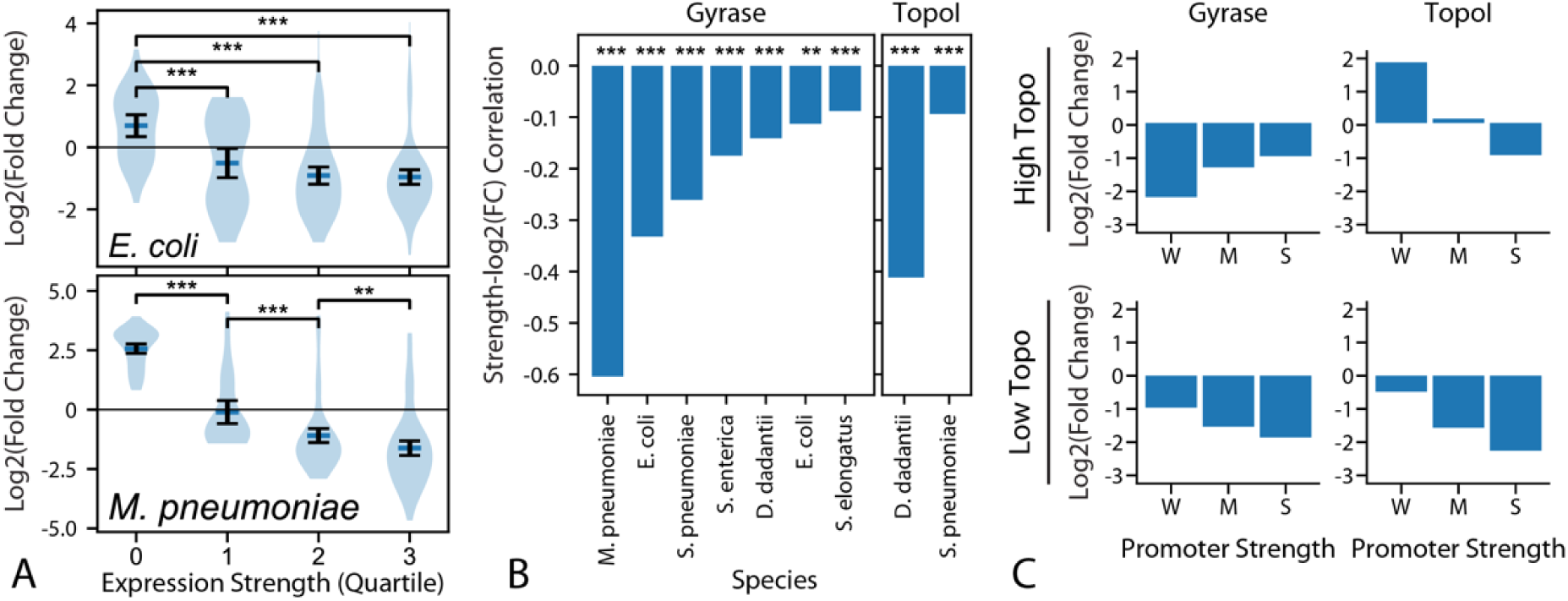
Transcriptome Response to Topoisomerase Inhibition. (A) Fold-change in expression of genes in *E. coli* and *M. pneumonia* in response to gyrase inhibition [9, 33]. Genes are split into quartiles based on expression strength. (B) Spearman’s correlation coefficient (and statistical significance) between gene expression strength and log2-fold-change in response to gyrase or TopoI inhibition in several bacterial species. The statistical significance is shown with symbols based on the correlation p-value: *** (*P* < 10^-3^) or ** (10^-3^ *< P* < 10^-2^). The list of conditions is detailed in S1 Table S1. (C) Log2-fold-change of simulated transcription rate in response to topoisomerase inhibition (5-fold reduction of either gyrase or TopoI activity). Top row: inhibition from high topoisomerase binding rates that are equal to the steady-state values (calculated to eliminate supercoils introduced by elongation, cyan lines in Figs. 3, 4 and 5B). (Bottom row) inhibition from low topoisomerase binding rates equal to 10% the steady state values.

To broaden these observations, we systematically analyzed the correlation between expression strength (without arbitrary classification into quartiles) and its response to gyrase or TopoI inhibition for 11 datasets obtained in 6 species (Fig. 6B). All analyzed datasets exhibit a clear negative correlation coefficient irrespective of gyrase or TopoI inhibition, indicating a statistically significant tendency towards repression of strongly expressed genes when gyrase or TopoI is inhibited. This robust observation is notable considering the strong heterogeneity of the analyzed studies. On the other hand, for several datasets, the correlation coefficient is relatively weak (but still significant due to the hundreds or thousands of genes considered), showing that the magnitude of this relation is probably quite dependent on the experimental conditions.

We then compared this trend to our simulation results. In the top row of figure 6C, we compare the fold-change in expression produced in weak (w, k_i_ = 0.008 s^-1^), moderate (m, k_i_ = 0.05 s^-1^), and strong (s, k_i_ = 0.2 s^-1^) genes of an intermediate distance to barriers (D = 10 kb) in response to a 5-fold reduction of either gyrase or TopoI activities from the steady-steady topoisomerase activities, which are sufficiently high to fully cancel the SC introduced by elongation. We see alignment between experiment and simulation for TopoI inhibition, which shows specific upregulation of weak genes, but not for gyrase, which shows the opposite trend in our model. However, when we apply the same 5-fold topoisomerase reduction to a gene being acted on by already reduced topoisomerase binding (10% the steady-state, Fig. 6C, bottom row), our model reproduces the experimentally observed relative upregulation of weak genes in response to both gyrase and TopoI inhibition. This observation matches with previous suggestions that most bacterial genes (thousands) are transcribed at a sufficiently low level to undergo sustained transcription with little local recruitment of topoisomerases, whereas the hundreds of active gyrase/TopoI molecules concentrate in a minority of topological domains with strong transcription (especially ribosomal operons) [50].

## Discussion

We have designed and implemented a model of supercoiling-coupled transcription allowing a systematic exploration of regulatory effects induced by topological constraints on a single gene. We used this model to directly compare with diverse experimental data, from *in vitro* transcription assays to *in vivo* genome-wide responses to topoisomerase inhibitions, as well as transcriptional bursting. The simulations replicated key observations from these datasets and thus provide a unifying framework to quantitatively model the transcription-supercoiling coupling. As such, this work is an important advancement from existing works in the field.

We have observed in our simulation that regulation by topoisomerases is dependent on promoter strength. While weak genes are always repressed by higher TopoI activities due to the relaxation of promoter DNA, strong genes require an intermediate level of TopoI activity to maximize transcription as too little TopoI activity results in RNAP stalling due to negative SC accumulation. Meanwhile, too much TopoI activity results in slowed initiation due to the removal of negative SC at the promoter. This differential regulation of weak and strong genes results from how RNAP stalling has a greater impact on the free promoter fraction for stronger genes. This is consistent with the observation that stronger genes are relatively repressed by TopoI inhibition across bacterial species [9–17, 33].

Our simulations show that the introduction of topological barriers near a gene has a more inhibitory effect in strong genes, and that the effects of this topological constraint are mediated by a reduction to the effective topoisomerase activities. This is consistent with a nonspecific binding mechanism *in-vivo*, as the introduction of topological constraints near a gene would result in the separation of the gene from nearby sites which topoisomerases can bind. Moreover, the observation that an inhibition of topoisomerase activities results in the preferential expression of weak genes is consistent with our analysis of transcriptomics data across the bacterial kingdom.

Our simulations also widely exhibit the experimentally observed phenomenon of transcriptional bursting, without requiring any external events of transcription factor binding/unbinding. Transcriptional bursts occur when the negative SC generated upstream of elongating RNAP molecules facilitates the binding of additional RNAP molecules, forming a DNA-embedded self-activating regulatory loop. This mechanism is dependent on the promoter obtaining an steady-state SC level that represses, but does not eliminate, initiation in the absence of already bound RNAP molecules. This Fano factor-maximizing SC level is more relaxed for strong promoters, which can exhibit extremely diverse behaviors, from sub-Poissonian to extremely bursty transcription, depending on topological conditions. Therefore, the topoisomerase binding rates that maximize the Fano factor are dependent on the promoter and topological domain length, suggesting that the behavior of a specific experimental system may not be easily controllable.

This observed behavior is largely consistent with observations in a wide range of promoters with varying expression strength and dependence on transcription factors. Fano factors in our simulations range from 0.66 to 4.75 (S4 Fig), consistent with observed values [28]. In the latter study, Chong et *al*. proposed that gyrase binding introduces gyrase-bound “on” states and gyrase-unbound “off” states, and that these binding-unbinding events are responsible for transcriptional bursting, as only the gyrase-bound states can relax the downstream positive SC introduced by RNAP elongation. Previous modeling work rather proposed that this mechanism results from extrinsic looping and unlooping stochastic events, inducing different topological constraints affecting transcription [23]. Here, we propose that even in the absence of an extrinsic regulatory factor, transcription within a topological domain can be intrinsically noisy, and that, under low gyrase activity, the on/off switching is controlled by RNAP binding dynamics, i.e., the strength of the promoter, rather than gyrase binding events. This alternative explanation is supported by the observation that Fano factors are only weakly affected by gyrase inhibition [28].

It is important to consider how our model compares to others in the field, in particular that used by Boulas et *al*. [17], one of few models compared to expression data. Boulas provides both experimental and computational evidence that transcription is mostly sensitive to the upstream (rather than downstream) barrier distance and that stronger promoters are more sensitive to that parameter. In this work, we varied upstream and downstream distances together and still observed that stronger genes are more strongly repressed by topological constraint, consistent with Boulas’ work. Our work provides a relation between this dataset and other sources of data, both *in vitro* and *in vivo*, and shows the regulatory effects of varying topoisomerase binding rates.

Several additional aspects would be interesting to consider. As mentioned previously, it has been shown that the relative arrangements of neighboring genes play a role in how genes are regulated by topoisomerase inhibition [13]. Thus, an expansion of our analysis to include multiple genes is justified. Our model is currently unable to replicate the cooperative effect of co-transcribing RNAPs with regards to elongation speed [51]. It is likely that reproducing this phenomenon would require refining our all-or-nothing response of elongation to SC, as already explored in other models [23–25], for example by introducing partial RNAP rotation during elongation. The ability of RNAPs to rotate varies based on the condition; for example, *in vivo* (with simultaneous translation) vs *in vitro;* RNAP transcribing genes for membrane-bound proteins are more constrained due to the insertion of the simultaneously translated peptide into the membrane [52]. Since the quantitative level of RNAP rotation remains largely unknown, it would be interesting to assess (either computationally or experimentally) how varied degrees of RNAP rotation influence the regulatory behavior of DNA topology.

## Roles

Conceptualization: Boaz Goldberg, Nicolás Yehya, Jie Xiao, Sam Meyer

Data Curation: Boaz Goldberg, Sam Meyer

Formal Analysis: Boaz Goldberg, Sam Meyer

Funding Acquisition: Jie Xiao, Sam Meyer

Investigation: Boaz Goldberg, Jie Xiao, Sam Meyer

Methodology: Boaz Goldberg, Nicolás Yehya, Jie Xiao, Sam Meyer

Project Administration: Jie Xiao, Sam Meyer

## Resources

Software: Boaz Goldberg

Supervision: Jie Xiao, Sam Meyer

## Validation

Visualization: Boaz Goldberg, Sam Meyer

Writing – Original Draft Preparation: Boaz Goldberg, Sam Meyer

Writing – Review & Editing: Boaz Goldberg, Nicolás Yehya, Jie Xiao, Sam Meyer

Corresponding Author: Sam Meyer, Jie Xiao

## Supporting Information

**S1 Table.**
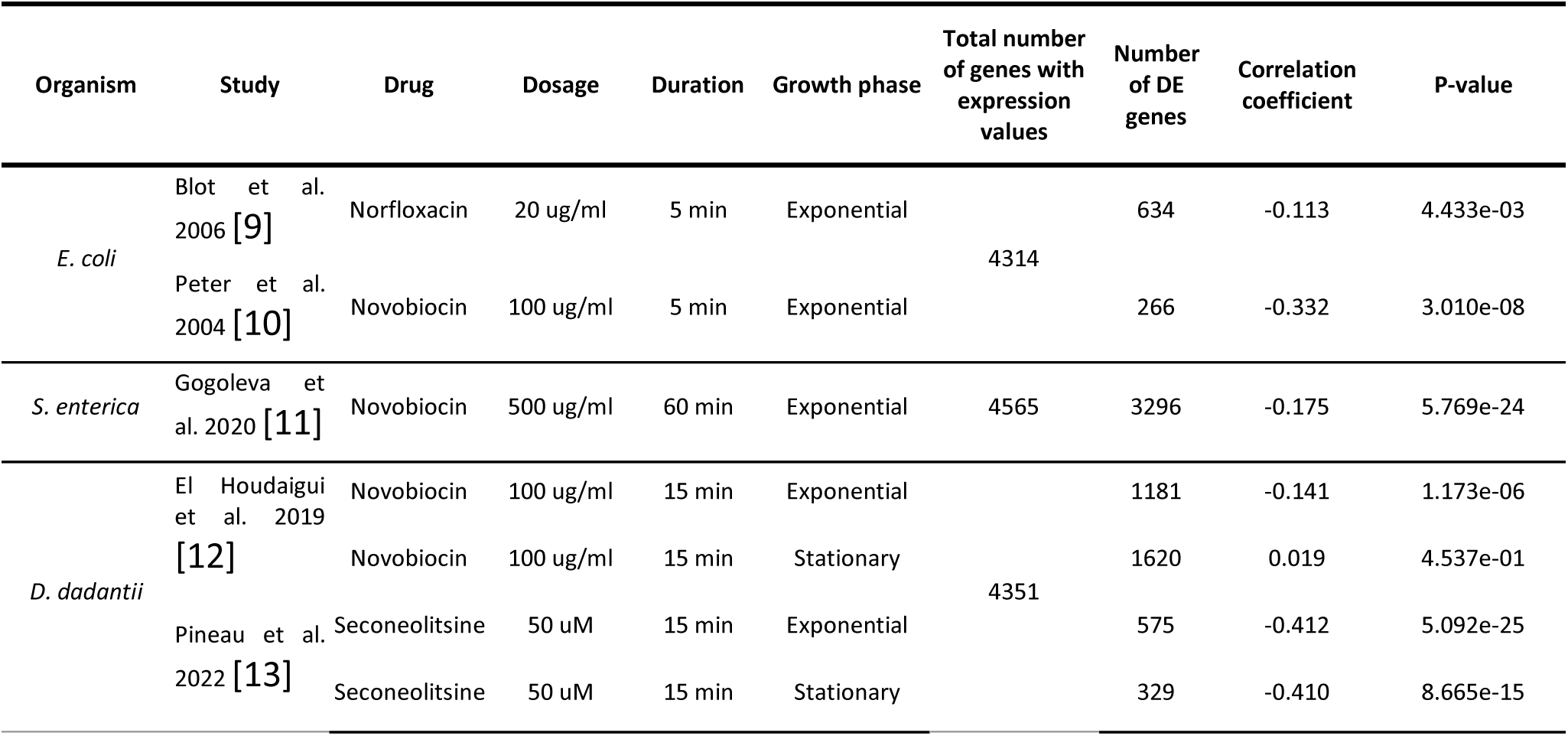
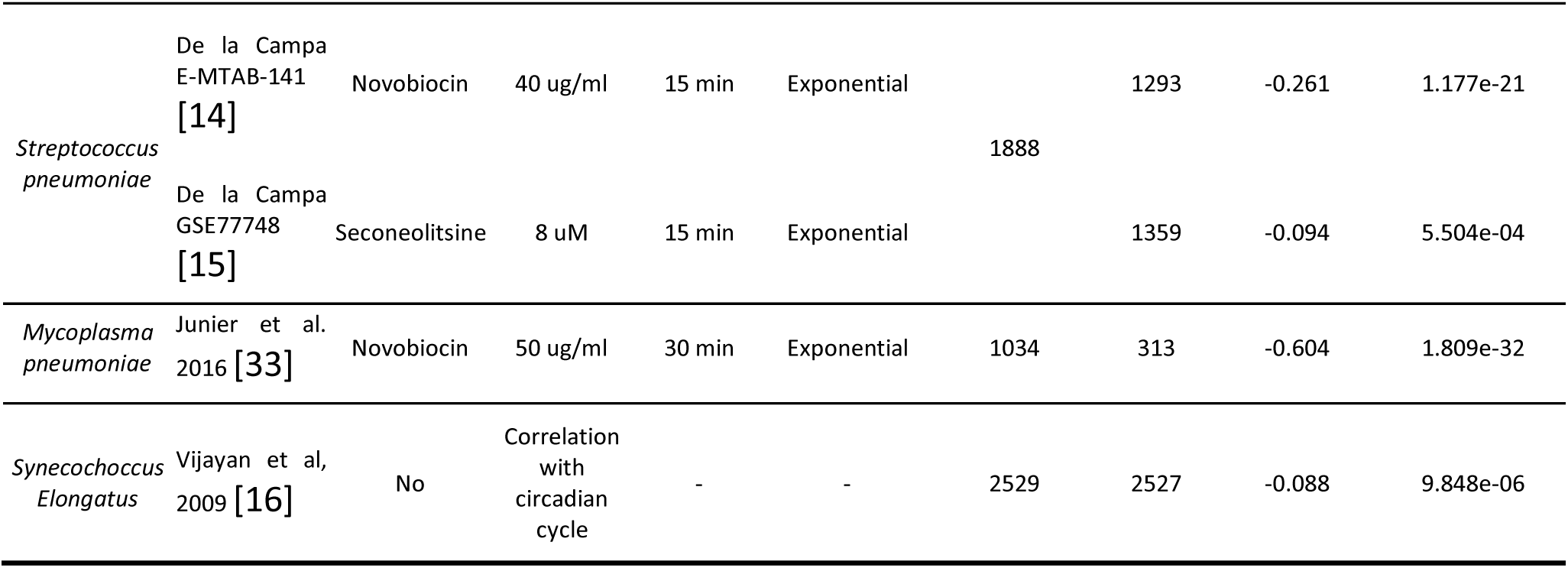
Sources and Conditions of Expression Data.

**S1 Fig.**
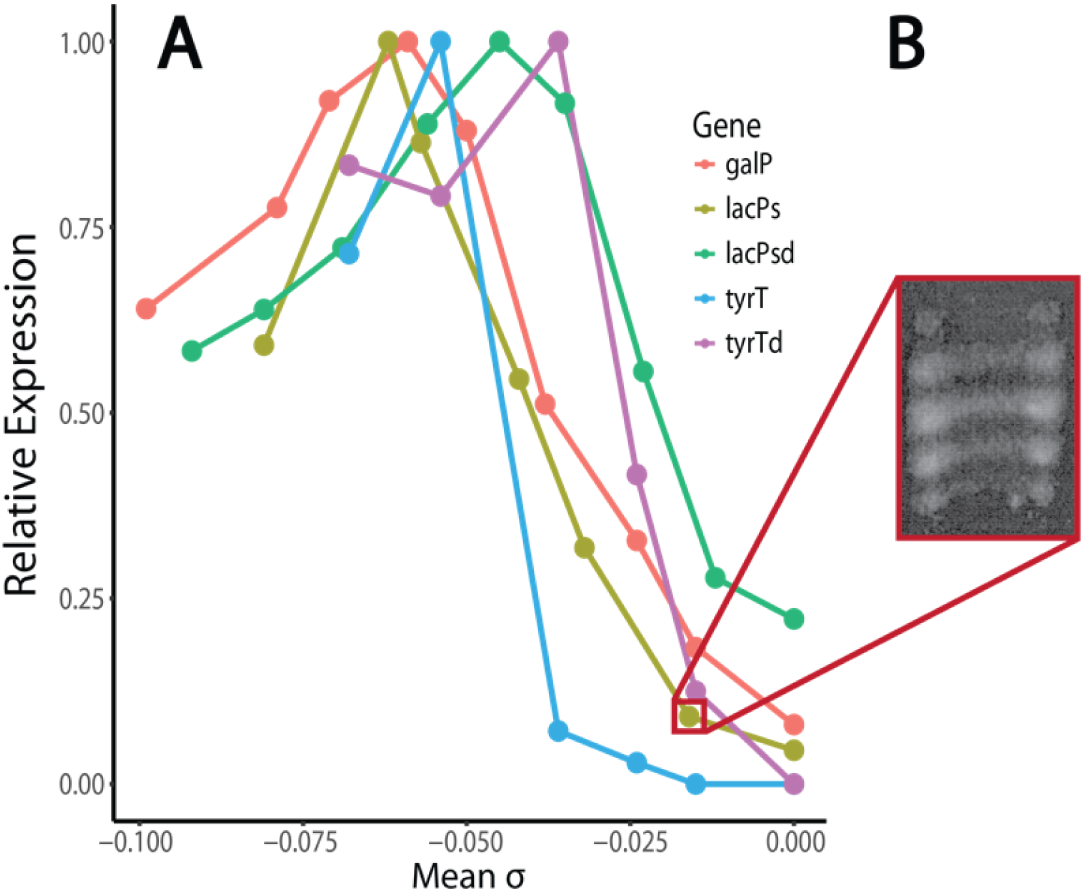
*In vitro* SC-dependence of transcription. (A) *In vitro* expression of several genes on plasmid samples of varied mean SC levels. The maximum expression rate for each gene is normalized to one [30–32]. (B) An illustrative gel to show the distribution of topoisomers within each sample [32].

**S2 Fig.**
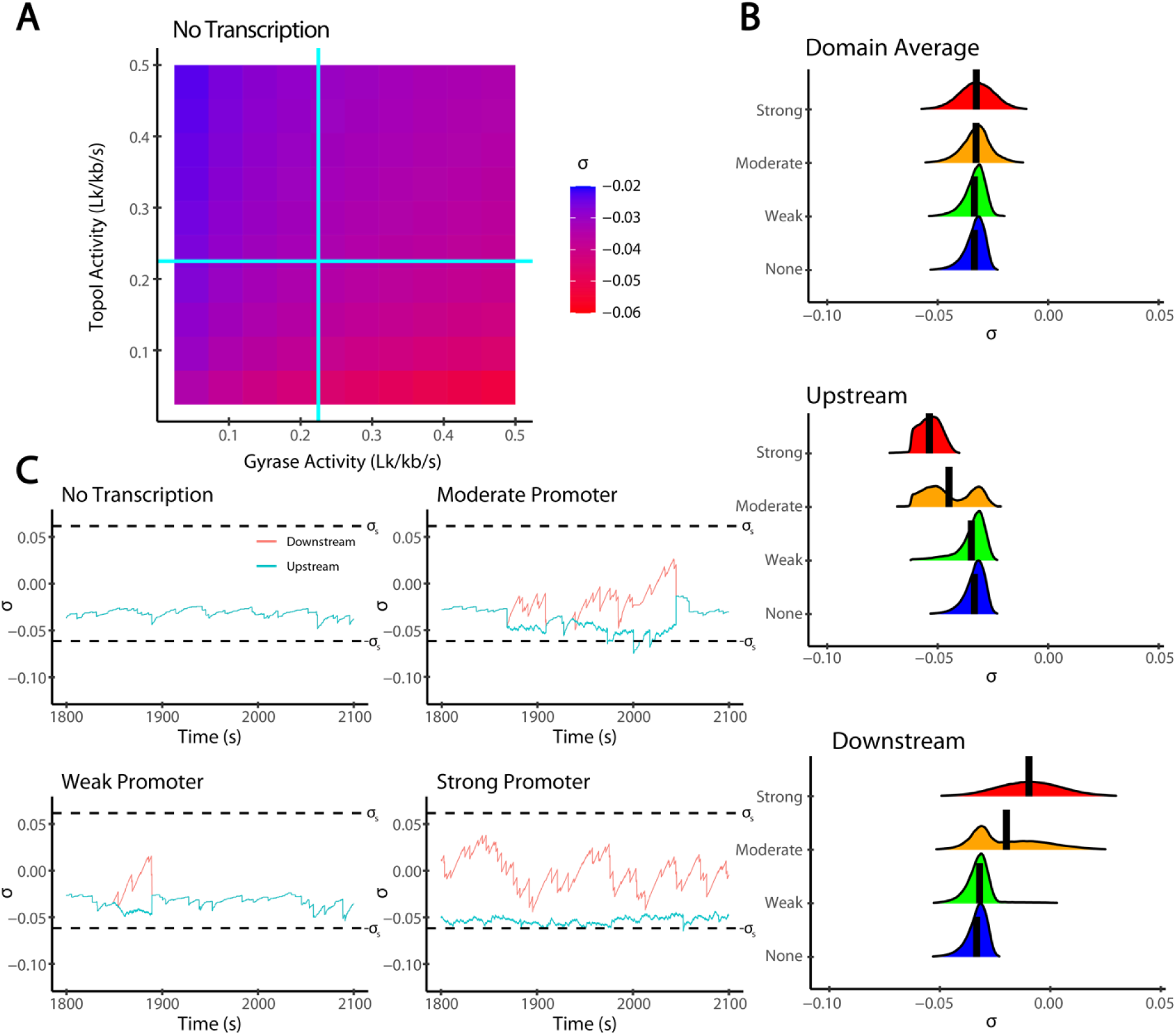
Impact of Topoisomerases and Transcription of Supercoiling. (A) Mean *σ* from simulations run in the absence of transcription at various topoisomerase binding rates. (B) Distribution of *σ* average across the domain, upstream of the gene, and downstream of the gene for simulations run at several promoter strengths (weak: *k*_i_ = 0.008 s^-1^, moderate: *k*_i_ = 0.05 s^-1^, strong: *k*_i_ = 0.2 s^-1^). Topoisomerase binding rates are the steady state values *k*_T_* and *k*_G_*. (C) Time courses of the upstream (blue) and downstream (red) *σ* values for several promoter strengths.

**S3 Fig.**
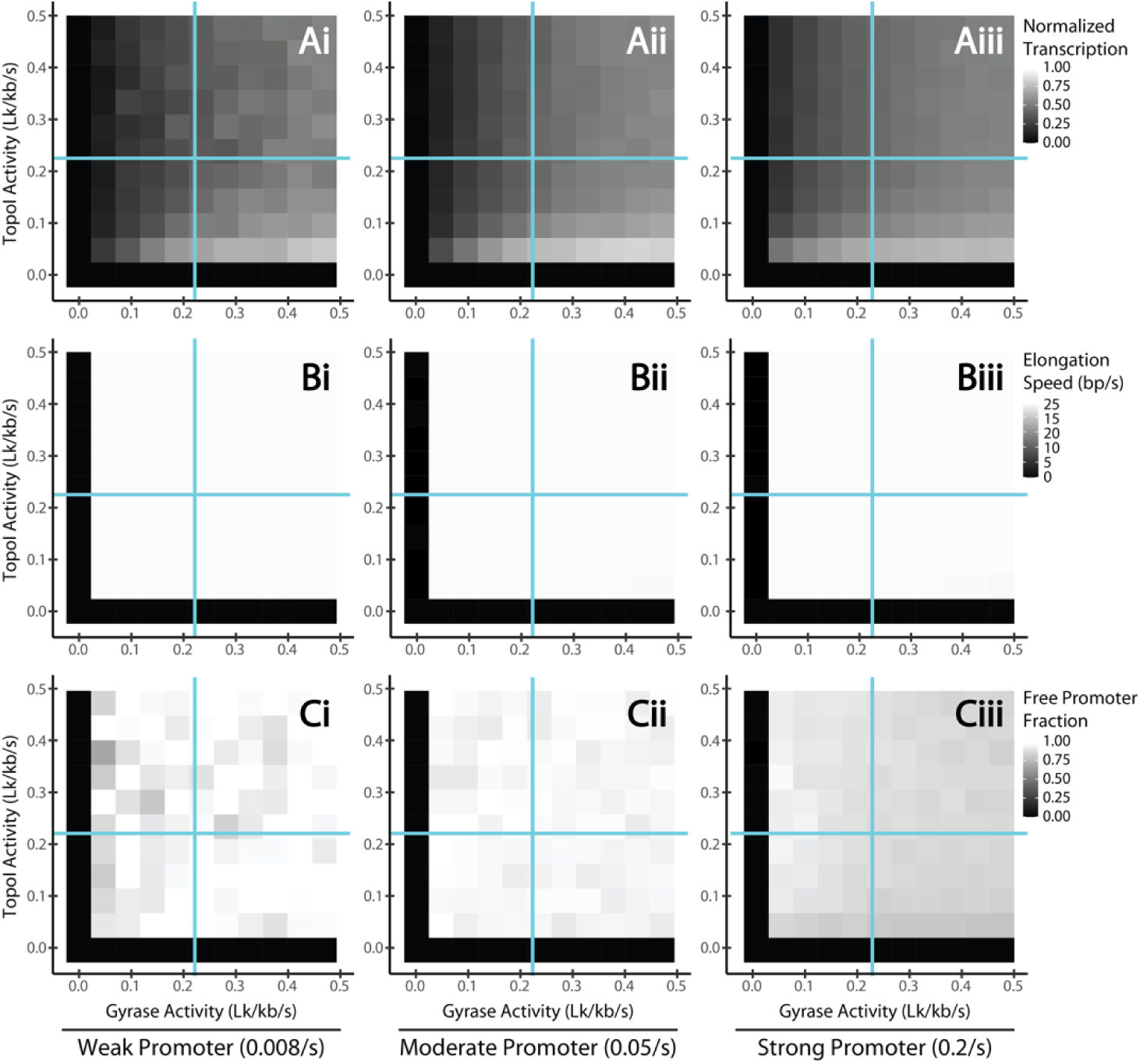
Regulatory effects of topoisomerases on transcription with distant topological barriers. Same as figure 3, replicated with distant topological barriers (*D* = 100 kb). Mean normalized transcription rate (each value is *k*_obs_/*k*_i_, i.e. normalized by the initiation rate of the promoter in the absence of supercoiling constraints, row A), mean elongation speed (bp/s, row B), and the free promoter fraction (the proportion of timesteps during which the promoter is not sterically occluded by a bound RNAP molecule, row C) are plotted as two-dimensional heatmaps at different combinations of Topo I (y-axis) and gyrase (x-axis) activities. We simulated 1 kb genes flanked by 100 kb barriers with three promoter strengths, weak, moderate and strong (columns i, ii, and iii respectively). The intersections of the two cyan lines denote the steady-state topoisomerase activities needed to cancel SC generated by elongation.

**S4 Fig.**
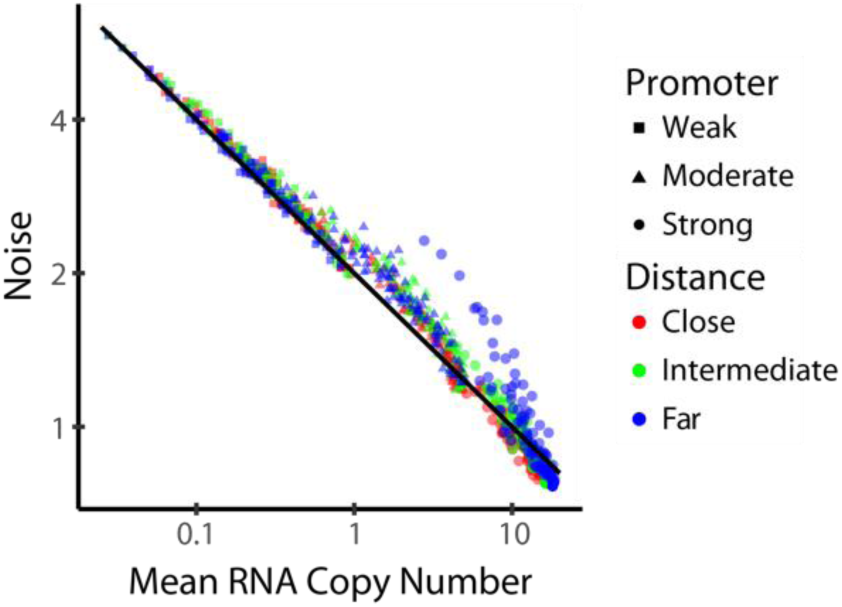
Noise and Expression Strength. The noise and mean RNA copy number for simulations ran with varied promoter strengths (weak: squares, *k*_i_ = 0.008 s^-1^, moderate: triangles, *k*_i_ = 0.05 s^-1^, strong: circles, *k*_i_ = 0.2 s^-1^) and barrier distances (close: red, *D* = 1 kb, intermediate: green, *D* = 10 kb, far: blue, *D* = 100 kb). The black line shows points where the noise is the reciprocal of the mean copy number, as expected for a Poisson distribution.

**S5 Fig.**
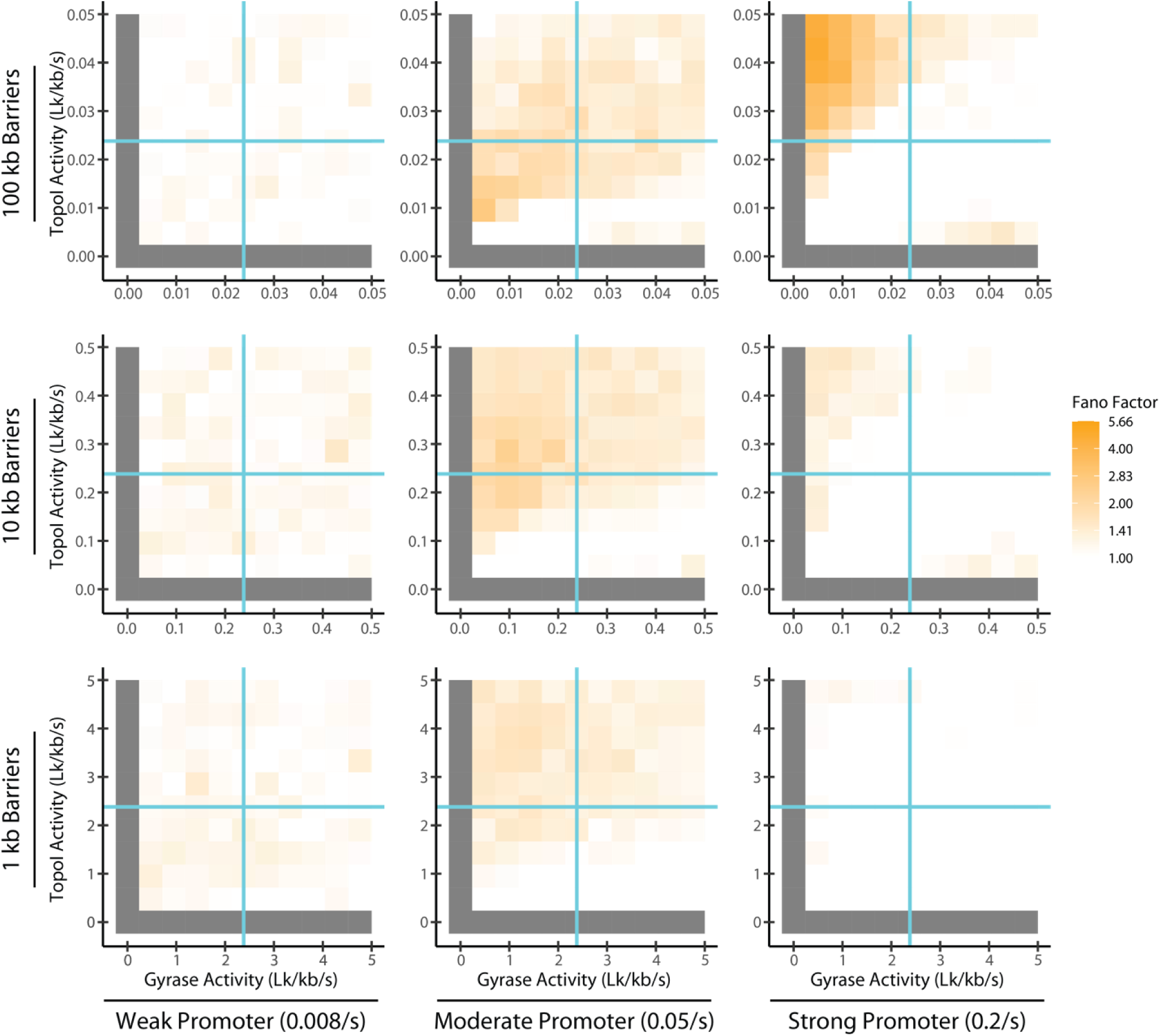
Simulated Fano Factors. Fano factors for simulations ran with varied promoter strengths, barrier distances, and topoisomerase activities. Note that topoisomerase activities were scaled such that the domain-wide topoisomerase binding rate (i.e., nonspecific binding rate times barrier distance) was constant across barrier conditions. Fano factors less than 1 are shown as 1. Such cases result from a forced spacing of initiation events due to bound RNAPs blocking the promoter. Fano factors are displayed on a logarithmic scale.

